# Cell signalling stabilizes morphogenesis against noise

**DOI:** 10.1101/590794

**Authors:** Pascal Hagolani, Roland Zimm, Miquel Marin-Riera, Isaac Salazar-Ciudad

**Affiliations:** Evo-devo Helsinki community, Centre of Excellence in Experimental and Computational Developmental Biology, Institute of Biotechnology, University of Helsinki, Helsinki, Finland; Institute of Functional Genomics, École Normale Superieure, France; European Molecular Biology Laboratory, Barcelona; Pompeu Fabra University, Barcelona; Genomics, Bioinformatics and Evolution. Departament de Genètica i Microbiologia, Universitat Autònoma de Barcelona, Cerdanyola del Vallès, Spain; Centre de Rercerca Matemàtica, Cerdanyola del Vallès, Spain

## Abstract

Embryonic development involves gene networks, extracellular signaling, cell behaviors (cell division, apoptosis, adhesion, etc.) and mechanical interactions. How should gene networks, extracellular signaling and cell behaviors be coordinated to lead to complex and robust morphologies?

To explore this question, we randomly wired genes and cell behaviors into a huge number of networks in EmbryoMaker. EmbryoMaker is a general mathematical model of animal development that simulates how embryos change, *i.e.* how the 3D spatial position of cells change, over time due such networks. Real gene networks are not random. Random networks, however, allow an unbiased view on the requirements for complex and robust development.

We found that the mere autonomous activation of cell behaviors, especially cell division and contraction, was able to lead to the development of complex morphologies. We also found that complex morphologies tend to be less robust to noise than simple morphologies. However, we found that morphologies that developed through extracellular signaling and complex gene networks were, on average, more robust to noise. This stabilization occurs when gene networks and extracellular signaling partition the embryo into different regions where cell behaviors are regulated in slightly different ways. Our results are consistent with theories proposing that morphological complexity arose in early metazoan evolution as a consequence of the cell bio-mechanics already present in protozoa and that robustness evolved by the co-option of gene networks and extracellular cell signaling.

## Introduction

There is no consensus definition of complexity, yet it is evident that organisms are complex and explaining such complexity is one of the most fundamental questions of biology. Morphological complexity is generated, in each generation, from a simple initial condition (*e.g.* a zygote) in a process called development. Morphological complexity also changes between generations in evolution. However, morphological complexity has not increased in the evolution of all lineages (Bonner, 2004; Williams, 1996; McCoy, 1977; Hinegardner & Engelberg, 1983; Gould, 2002; Arendt, 2008; Canning & Okamura, 2004; Arthur, 2011) and, in general, it is unclear whether there is a general trend of increasing complexity in evolution (Fisher, 1986; Ruse, 2009; Gould, 2002; McShea, 1996), but see (Fleming & Mcshea, 2013). Yet, one may ask about the mechanisms by which such complexity has increased in the lineages where it has increased.

How complexity increases during evolution is necessarily related to development: any evolutionary change in morphology is first a change in the development that produces such morphology. The process of development can be described as a sequence of transformations of specific distributions of cell types in space, what we call a developmental pattern, into other, usually more complex, developmental patterns (Salazar-Ciudad, 2003). The first such developmental patterns would be, for example, the zygote while the last would be the adult phenotype.

It has been argued that, in spite of the remarkable complexity of organisms, their development is achieved through a limited number of cell behaviors and types of cell interactions (Salazar-Ciudad, 2003; Davies, 2013; Newman & Bhat, 2009). These cell behaviors would be cell division, cell adhesion, cell death, cell growth, cell contraction, extracellular signal and matrix secretion, extracellular signal reception and cell differentiation (Salazar-Ciudad, 2003; Davies, 2013; Newman & Bhat, 2009). One may consider, in addition, cell migration and cell shape changes as resulting from specific patterns of cell contraction and adhesion.

The two main types of cell interactions in development are cell signaling, and cell mechanical interactions arising from forces generated by cell behaviors (*e.g.* cell contraction). Cell signaling typically occurs through the secretion of extracellular diffusible molecules by some cells and the reception of those by other cells but it can also occur through membrane-bound signals or by other ways (Gilbert and Barresi, 2016). Both types of interactions can lead to gene expression changes and those to changes in the behaviors of cells (Gilbert and Barresi, 2016). Which cells express which genes, however, is also affected by cell behaviors since these affect the distribution of cells in space and this, in turn, affects the distribution of extracellular signals and forces in space (Salazar-Ciudad, 2003). In this article, as in (Salazar-Ciudad, 2003), we define a *developmental mechanism* as a network of gene interactions, cell interactions and cell behaviors required for the transformation of one developmental pattern into another, see supplementary figure S1. The gene network part of a developmental mechanism is not a specification of whether two gene products actually interact in a given cell during development but of, whether and how, these two gene products would interact if they happen to coincide in a cell. Where and when this may happen is determined by the dynamics of each developmental mechanism and of the whole of development.

The question we want to approach in this study is: how should these interactions and cell behaviors be coordinated to produce complex and robust morphologies? The question is, then, whether there are some logical requirements that developmental mechanisms should fulfill in order to lead to complex robust morphologies. Are there, for example, some requirements at the level of gene network topology or at the level of cell behaviors and their coordination during development?

A large proportion of the literature in developmental biology provides insights into the development of complex morphology for specific organs. Some literature also exist on the robustness of development in some organs (Barkoulas *et al*, 2013; Oliveira *et al*, 2014). Here, instead, we use a mathematical model of development to try to address this question in general. Our model is, as any model, a simplification of nature, but it is not constrained by the specificities of any developmental system and, thus, our conclusions should apply to the development of any animal, as we explain in the following lines.

If, as suggested above, pattern transformations in development involve a limited set of cell behaviors and cell interactions, then any mathematical model implementing those and intracellular gene networks should be able to reproduce, to a large extent, the range of pattern transformations possible in animal development. In this work we use one such model, EmbryoMaker (Marin-Riera *et al*, 2015), to simulate many different developmental mechanisms and try to discover what, if anything, do the mechanisms leading to robust complex morphologies have in common. To that end, we randomly wired genes and cell behaviors into developmental mechanisms and simulated, with EmbryoMaker, the morphologies they produce from a simple initial developmental pattern. This initial condition consist of a small flat epithelium with an underlying layer of mesenchymal cells (see supplementary figure S1 and S2). Real developmental mechanisms are not random but the result of millions of years of evolution. However, the study of random developmental mechanisms allows us to identify general requirements without being conditioned by our current understanding of development. In spite of its statistical nature, our approach should also be informative about evolution since the developmental mechanisms found in current animals may still need to fulfill these general requirements.

EmbryoMaker is a general mathematical model of animal development in the sense that it can simulate all the basic behaviors of animal cells, extracellular signal-mediated interactions and mechanical interactions between cells as well as any arbitrary intracellular gene network (see Figure 1). EmbryoMaker represents cells as a set of parts (herein called nodes) with specific mechanical properties. Mesenchymal cells are represented by single spherical nodes and epithelial cells by a cylinder consisting of two nodes (one basal and one apical bound by an elastic link). Nodes touching each other experience adhesion forces but, if they get closer than a given distance, they experience a repulsive force (supplementary figure S3). Epithelial cells exert additional forces into neighboring epithelial nodes that reflect their specific mechanical properties and their organization in epithelial sheets (supplementary figure S3 and SI 1.2).

**Figure 1.**
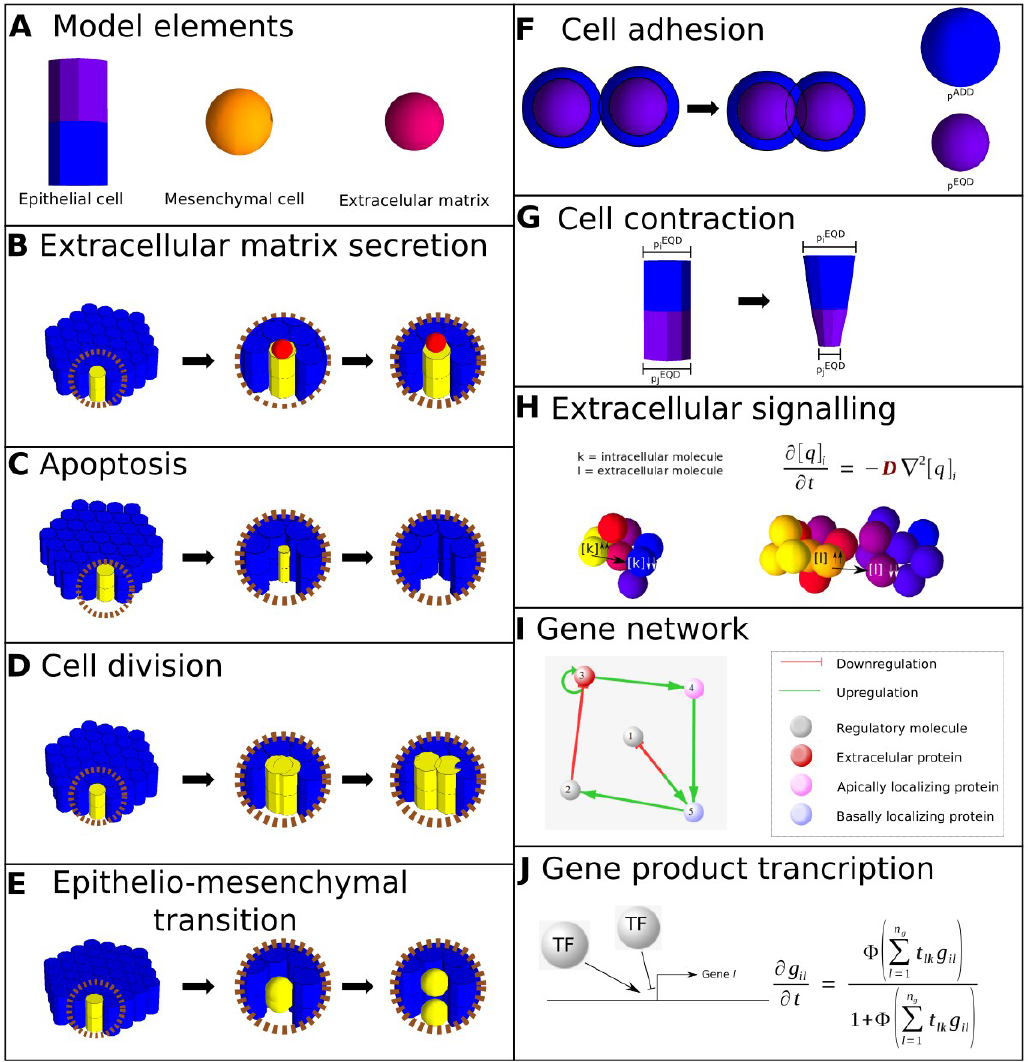
EmbryoMaker. **A.** EmbryoMaker includes three types of elements or nodes. Epithelial cells (cylinder) are made of two nodes, the basal one in blue and the apical one in violet. Mesenchymal cells and extracellular matrix elements are made of single spherical nodes. **B-H** Cell behaviors used in EmbryoMaker. **B.** Extracelular matrix secretion. Any cell type can secrete ECM, given that a there is expression of a molecule regulating its secretion. **C.** Apoptosis. When a cell undergoes apoptosis, its size will decrease until it reaches a minimal value, at which point it is completely eliminated. **D.** Cell division. The plane of division of the cells is normal to the longest axis of the cell. **E.** Epithelial-mesenchymal transition. An epithelial cell is transformed into a mesenchymal cell by turning the cylinder into a pair of spheric nodes. **F.** Cell adhesion. Two cells that are in contact, i.e., their radius of adhesion overlap (*p*^*ADD*^, blue sphere) will come closer to each other until their equilibrium distance is reached (*p*^*EQD*^, purple sphere). **G.** Cell contraction. Parts of the cell can change their size by decreasing the equilibrium radius. If this happens in one of the nodes of an epithelial cell, the cylinder will change its shape to a more conic one. **H.** Extracellular signaling. Diffusion is implemented as transfers of molecules between nodes. This transports follows Fick’s second law of diffusion (see top-right side). Letters in red denote parameters of the model while letters in black denote variables. **I.** Example of the gene network of a developmental mechanism. The spheres depict gene products and the arrows transcriptional regulation. There are also several types of gene products: extracelular, they diffuse extracellularly; apically and basally locates gene products. **J.** Gene product transcriptional interactions. *g*_*il*_ is the amount of transcriptional factor *l* in node *i* and each *t*_*lk*_ term is the strength by which each specific transcriptional factor *k* activates or inhibits the transcription if gene *l*.

Within each node there are gene products that can affect the expression of other gene products and the mechanical properties and cell behaviors of their containing nodes (*e.g.* their size, distance at which repulsion forces apply, adhesion, mitosis rates, elasticity, etc.). Some of these molecules can diffuse in extracellular space and affect cells other than the cell where they are produced, thus leading to extracellular signaling. Nodes’ mechanical properties determine how they respond to forces (for example node size, node’s elasticity). At the mathematical level the concentration of each molecule in a node, a node’s mechanical properties and a node’s 3D position are continuous variables that are calculated by differential equations that take into account some of these same variables in neighboring nodes (see methods and (Marin-Riera *et al*, 2015) for details).

EmbryoMaker is not a model of the development of any specific system. Models for specific systems *e.g.* a model of limb development, are built in EmbryoMaker by specifying a concrete developmental mechanism: which gene products regulate other gene products, mechanical properties and cell behaviors in a specific system. As a result of an initial developmental pattern and a developmental mechanism, EmbryoMaker simulates development and outputs how the spatial distribution of cells and gene expression in the initial pattern change over time until some final developmental pattern (a morphology) is reached (see supplementary figure S1). On these virtual morphologies we measure complexity and robustness.

There is no consensus definition of complexity and any study using one is likely to be controversial. Because of that, we use two different quantitative measures of morphological complexity and stress that our results do not necessarily apply to complexity in general but to complexity as defined by these measures. There are other possible measures of complexity (Saunders, Work and Nikolaeva, 1999; McShea, 1996; Fleming and Mcshea, 2013) but they are not as easy to quantify in an objective way as we attempt in here or are not easy to directly apply to 3D morphologies. Roughly, our two measures reflect the likelihood of randomly guessing the position of a cell in 3D space knowing the position of its neighbors at different distances but knowing nothing about the developmental mechanism that produced such morphology. The first measure of complexity we use is angle variance, or AV: the variance of the angles between the polarity axis of epithelial cells at different distance intervals (see methods for a full description and figure S4 for examples). The second measure we use is the so called *orientation patch count* or OPC (Evans et al. 2007), the number of regions with the same slope orientation (see methods for a full description and figure S4 for examples).

Robustness is also a concept that is defined and understood in many different ways in the literature (Arjan *et al*, 2003; Salazar-Ciudad, 2007). Here we restrict ourselves to robustness understood as the suppression of developmental instability (Shapiro, 1971; Waddington, 1942; Klingenberg, 2002). For the rest of this article we will only talk about developmental instability while acknowledging that there are many other meanings to robustness that are not related to developmental instability (Salazar-Ciudad, 2007). This is how different are the morphologies of genotypically identical individuals that develop in exactly the same environment. This is morphological variation arising from noise in the developmental process itself. To measure developmental instability each developmental mechanism is simulated several times from the same initial developmental pattern but with noise. Noise is implemented as small random displacements of cells’ positions in each iteration of the model. The morphological distance between the resulting embryos is then a measure of developmental instability. We use two different complementary measures of morphological distance (see methods and SI 3.0 and Figure S5 for examples).

## Results

We built 20,000 random developmental mechanisms and run them in four different ways. In the signaling *ensemble* an initial developmental pattern with a gene expressed in a gradient was used (see methods and Figure 2). In the *gradient autonomous ensemble* we preclude extracellular signaling. In the *autonomous ensemble*, there is no extracellular signaling and one gene is homogeneously expressed through all cells while the rest of genes are not expressed at all. In the *autonomous bio-mechanics-only ensemble* there are only three non-interacting genes and these are expressed everywhere in the initial developmental pattern. These three autonomous ensembles are controls showing the morphogenesis that is possible from the uniform or gradient regulation of cell behaviors and mechanical properties, no extracellular signaling and, for *autonomous bio-mechanics-only ensemble,* no changes in gene expression over time.

**Figure 2.**
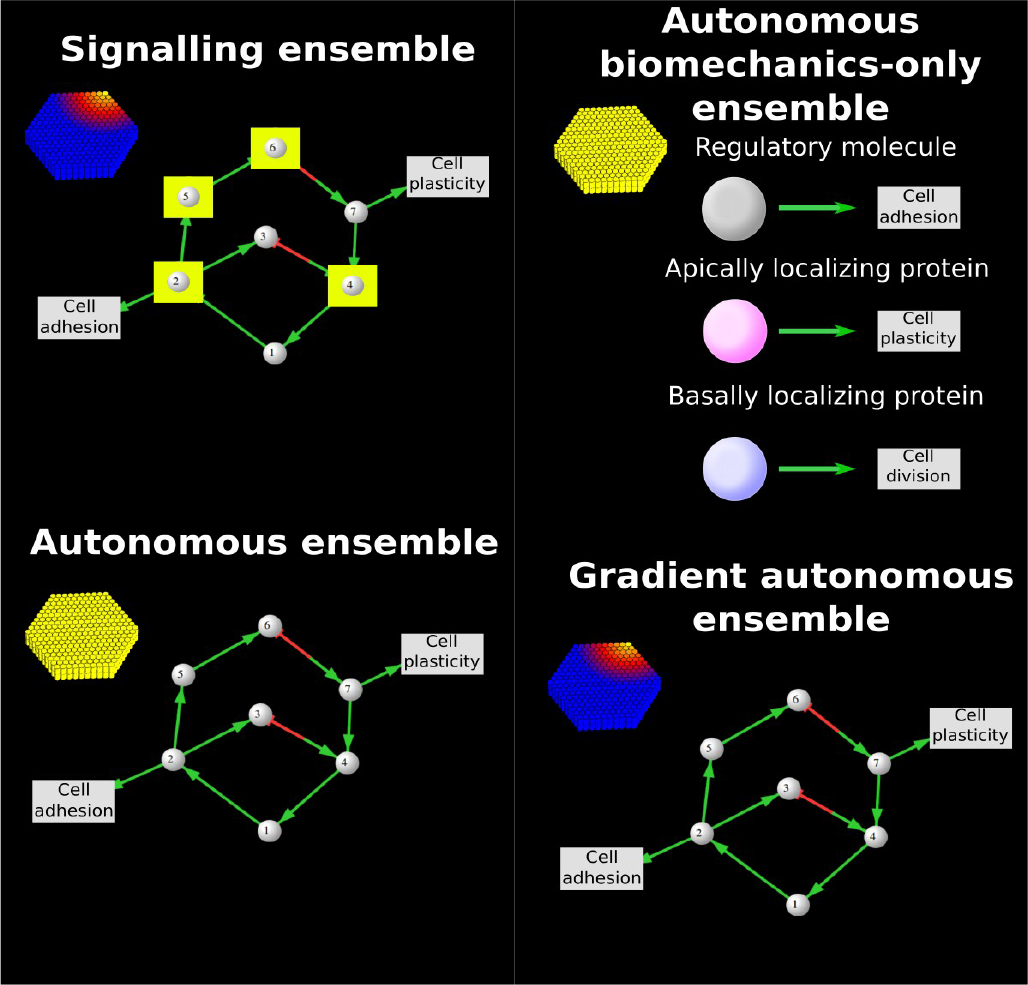
The ensembles. The initial developmental pattern of each ensemble is shown in the upper left side for each ensemble. In all ensembles such pattern started as a flat hexagonal embryo. In color we show the level of expression of the gene expressed in the initial condition (yellow is the maximal expression over the embryo and blue is the minimal). Next to the initial conditions we depict idealized examples of developmental mechanisms for each ensemble. The developmental mechanisms are represented as in figure supplementary figure S1. Circles represent gene products and arrows regulatory interactions (positive in green and negative in red) either on other gene products or on cell behaviors or node mechanical properties (these latter are shown in gray boxes). The circles surrounded by yellow boxes indicate growth factors (*i.e.* extracellular signaling).

In all ensembles we found complex morphologies, although at a low frequency (see figure 3 and figure S8 and supplementary figure S9 for some example morphologies). In addition, we found that the developmental mechanisms producing complex morphologies tend to be more developmentally unstable than the developmental mechanisms producing simple morphologies (see figure S8).

**Figure 3.**
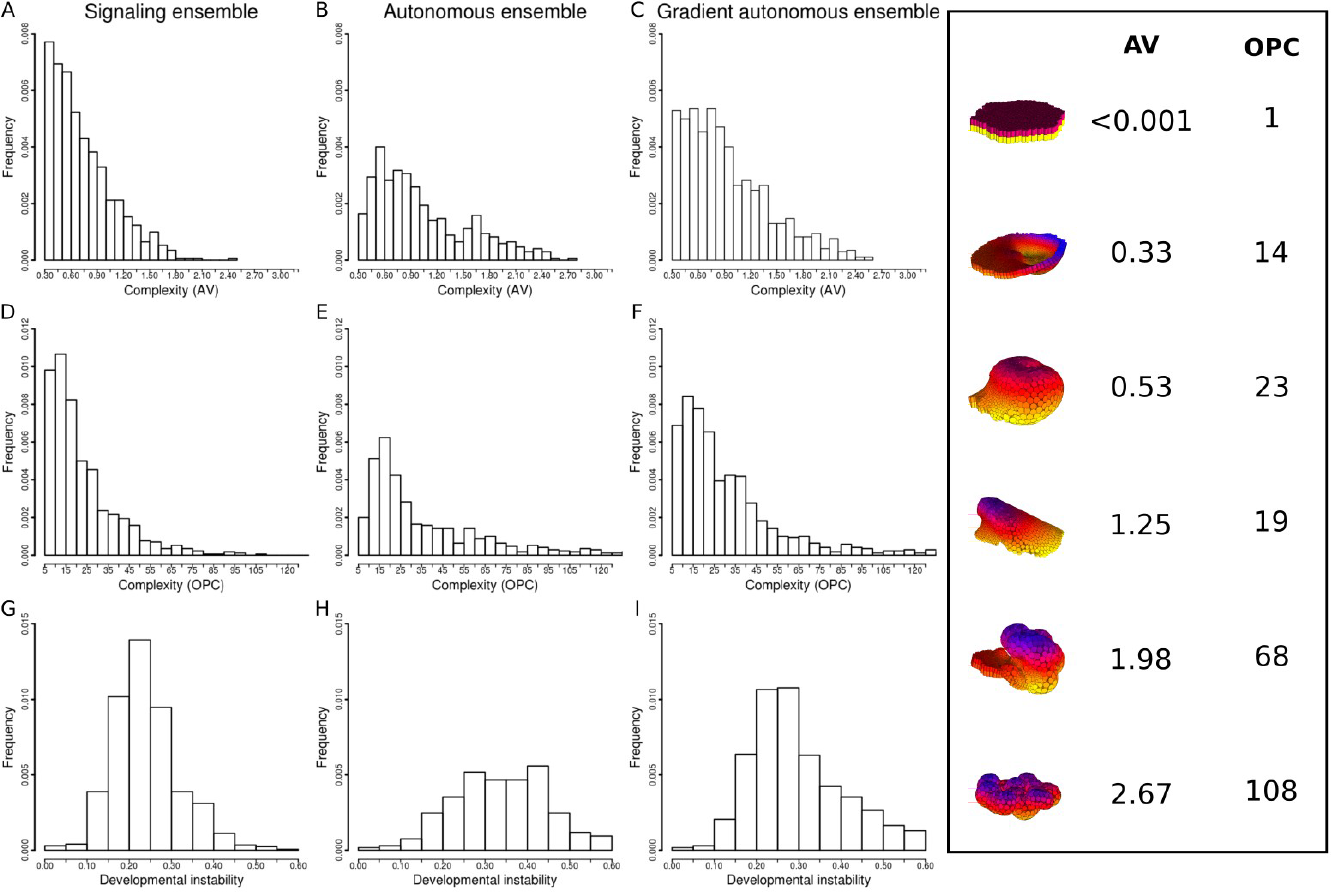
Lower complexity but higher stability is found in the signaling ensemble. The first column (A, D, G) shows the results for the signaling ensemble, the second column (B, E, H) for the autonomous ensemble and the third column (C, F, I) for the gradient autonomous ensemble. Upper plots show the morphological complexity as measured by AV (X-axis) versus proportion of the morphologies in each interval of complexity (Y-axis) for the three different ensembles. Middle plots: the same as the upper plots but for OPC complexity. Lower plots: Developmental instability (X-axis) versus proportion of the morphologies in the ensemble within each interval of complexity (Y-axis). The proportions are calculated based on the total size of each if the ensembles. The total size of each ensemble is 20,000. To facilitate visualization, all morphologies with an AV smaller than 0.3 are not considered in the upper and lower rows. For the OPC histograms (middle row), morphologies with an OPC smaller than 5 are not considered either. All the distributions are significantly different according a Mann-Whitney test. A to B distributions: p-val <0.001, Z=16.146, n=1340. A to C distributions: p-val <0.001, Z=10.701, n=1647. B to C distributions: p-val<0.001, Z=7.566, n=1391. D to E distributions: p-val <0.001, Z=10.701, n=1444. D to F p-val<0.001, Z=7.849, n=1786. E to F distributions: p-val<0.001, Z=3.843, n=1538. G to H distributions: p-val <0.001, Z=10.495, n=1387. G to I p-val<0.001, Z=8.612, n=1721. H to I distributions: p-val<0.001, Z=2.944, n=1486. The images on the right show example morphologies that can be found in the bins of complexity.

Most remarkably we found that the signaling ensemble produced morphologies that were, on average, significantly less complex than the morphologies produced in the two autonomous ensembles (see figure 3). The within-ensemble disparity was slightly smaller in the signaling ensemble than in the autonomous ensembles (see figure S10). Disparity was measured as the sum of the morphological distances between each morphology in an ensemble and all other morphologies in the same ensemble divided by the number of distances measured.

The four ensembles, however, differed in developmental instability (see figure 3 and figure S11). For morphologies of the same complexity, more developmentally stable morphologies were found in the signaling ensemble and in the gradient autonomous ensemble than in the two other autonomous ensembles (see figure S8). In other words, while extracellular signaling or gradients did not seem to be required for complex morphologies, they were required for complex morphologies to be developmentally stable.

Most complex morphologies in the autonomous and autonomous bio-mechanics-only ensembles consisted of highly folded epithelia, like crumpled paper balls, in which the position of epithelial folds was different in each run of the same developmental mechanism. In the two other ensembles, many morphologies were also composed, totally or partially, of randomly folded epithelia but there were also complex morphologies made of folds that consistently appeared in the same location and, thus, were complex yet developmentally stable (see supplementary figure S9 for examples of both kinds of morphologies).

We found that the development of complex morphologies requires only that, over developmental time, a large proportion of the cells in an embryo change the regulation of cell behaviors or mechanical properties (figure S12). Cell contraction and cell division were the cell behaviors most often associated with the development of complex morphologies (see figure S13). This is especially the case, but not exclusively, when cell contraction is asymmetric between the apical and basal side of each epithelial cell, as found in the formation of invagination and tubes in many animals (Martin & Goldstein, 2014). Cell division could lead to some buckling and wrinkling of the epithelium and then to some complexity, as in fact also observed in the development of many animal organs (Bunn *et al*, 2011; Striedter *et al*, 2015). This was specially the case if the epithelium had regions with different values in nodes’ mechanical properties or different rates of cell division. The regulation of other mechanical properties and cell behaviors also had an effect on complexity but to a lesser extent and only when contraction was also present (supplementary figure S14 and figure S15).

To explain how extracellular signaling enhances robustness to small noise affecting the movement of cells, we first paid attention to the effect of noise on development when there is cell contraction since this is the cell behavior most often associated with complex morphologies in our simulations. Qualitatively we found that, if cells contract at the same time and with the same intensity over large regions of the embryo, the resulting morphologies tend to be complex but unstable. If, on the contrary, cell contraction, occurs in different ways (at different moments or at different rates) over different regions of the embryo, development tends to lead to more complex but also more stable morphologies.

On purely geometric grounds it can be seen, see supplementary figure S16, that only a very specific number of cells can fit into an invagination. The larger cell contraction, *i.e.* the larger is apical side versus basal side or *vice versa*, the smaller is such number and the higher the curvature of the resulting invagination. This implies that, in our model, large fields of contracting cells will necessarily split into different invaginations and we found, that in that case, the positioning of such multiple invaginations is quite sensitive to noise.

By following the development of many complex multi-invaginated morphologies in the ensembles, we found that the positioning of invaginations was highly dependent on noise and that the instability to noise was related to the non-linear nature of the invagination process. As an invagination starts to form in our *in silico* embryos, its cells start to rotate its longest axis to align it with that of its also rotating neighbors. The rotation is, thus, in the direction in which neighbor epithelial cells have already contracted and rotated the most. In turn, the direction in which a cell rotates also affects the direction in which its neighbor cells rotate, further strengthening the rotation in one direction or another. This non-linear interdependence makes that, when invaginations start to form, slight noise in the timing, rate or direction in which cells contract, gets easily amplified over time and affects where each invagination will form.

To explore the quantitative support of these qualitative observations, we ran several simulations in which epithelia of different sizes contracted all their cells in the same way and at the same time (as in the cell behaviors-only ensemble). As it can be seen in figure 4 and supplementary figure S17 the larger the epithelium, the larger is its developmental instability. Splitting the epithelium into regions contracting in slightly different moments or at slightly different rates, however, decreased developmental instability (figure S18). The same occurred if the epithelium was split into equally contracting regions separated by narrow non-contracting boundaries (Figure S18). In other words, partitioning the embryo in different regions largely reduces developmental instability without precluding the development of complex morphologies.

**Figure 4.**
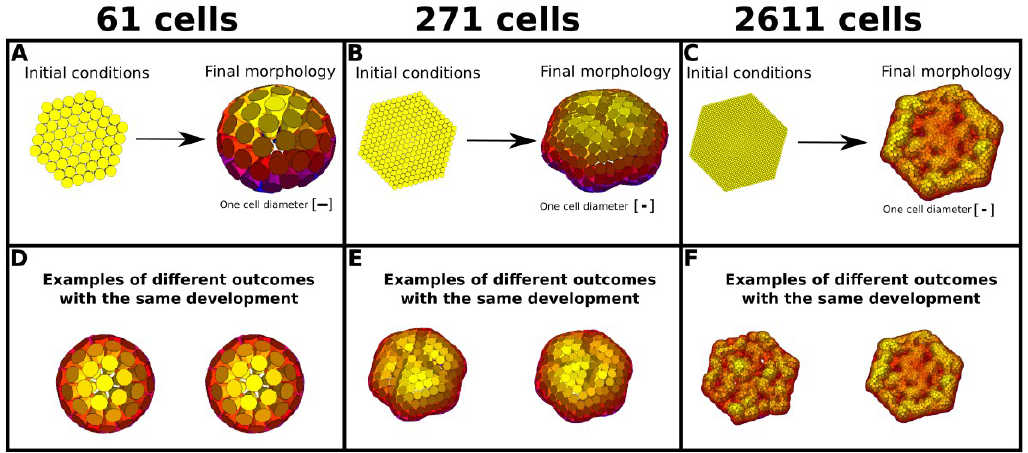
The larger the number of cells contracting at the same time the larger the developmental instability. Three different simulations where cell contraction occurs homogeneously over the epithelium are shown. A, B and C differ in the number of cells developmental patterns are shown. D, E and F depict two different runs from the same initial conditions. As it can be seen the arising morphologies are more different from each other when the initial developmental pattern includes more cells.

If developmental instability is related to the size of the epithelial regions contracting in the same way, then developmental instability and such size should correlate in the ensembles we simulated. Figure 5 shows that this was indeed the case: the developmental stability of an embryo correlates with the size (in number of cells) of the largest regions in which cells are contracting in the same way (*i.e.* they change their apical or basal side at the same rate). These regions are larger in the autonomous ensemble and cell behaviors-only ensemble since, in such ensembles, all cells behave in exactly the same way. All the embryos in figure 5 (and most embryos in the ensembles) have the same number of cells so the relationship we found was not due to larger embryos being less stable.

**Figure 5.**
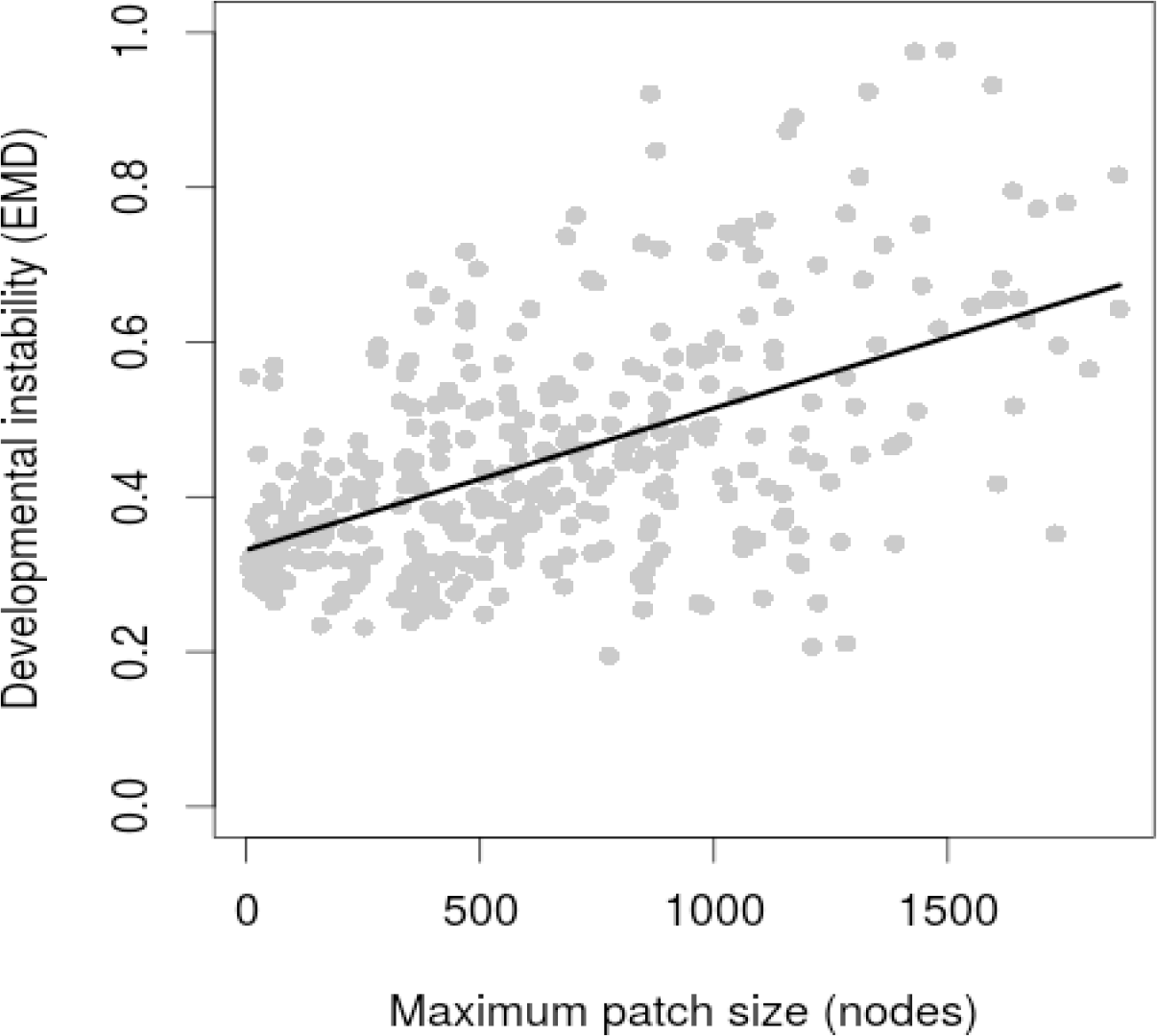
The number of cells contracting in the same way in an embryo correlates with developmental instability. For all morphologies in the signaling ensemble we calculated the number of cells that have a similar radius and are in contact to each other as forming a single patch (six groups are made, from 0.1 to 0.5 in 0.1 intervals). This was done for epithelial cells, taking into account apical and basal nodes separately. Then we calculate the size of the largest of such patches in each morphology. For each embryo in the signaling ensemble the maximal patch size is plotted against developmental instability. Only morphologies with a total size of at least 1000 epithelial nodes that have at least two patches and whose complexity by angle variation complexity was higher than 0.3 were included. The black line shows the lineal regression. Spearman correlation: r_s_=0.493, pval<0.001, n=440. Pval of correlation calculated with a permutation test.

The stabilizing effect of extracellular signaling was not found to be related to cell division. In contrast with contraction, partitioning the embryo into regions where cell division would occur at different moments or in slightly different ways, as done for contraction in Figure S18, had no clear effect on developmental instability (see Figure S19).

## Discussion

The main question of this article was whether there are some features of developmental mechanisms, *e.g.* at the level of gene network topology or cell behaviors, that are required for the development of complex robust morphologies. For the case of gene network topology we found the answer to be “no”. The fact that complex morphologies were less common in the signaling ensemble than in the ensembles without signaling suggests that the development of morphological complexity as such does not require extracellular signaling, at least for the range of complexity observed in our study. This suggest that the extracellular signaling we observe in current animals, or even the genetic regulation of cell behaviors over time, may not have arisen to build complex morphology *per se*, but rather, as we later detail, to make such complexity developmentally stable.

The answer to the main question of this article for the case of cell behaviors is “yes”, there is a simple requirement at the level of cell behaviors for the development of complex morphologies: the more cells in an embryo activate cell behaviors, the larger is, on average, the complexity of the arising morphology. This was specially the case for cell contraction and division. That complex morphology is often associated with cell division and contraction and that the other cell behaviors have only a modifying effect should not be surprising. In epithelia, these are the cell behaviors that can generate forces (Bard, 1992) and, thus, cell movement leading to morphogenesis. The secretion of extracellular matrix can also generate forces but we rarely found such cell behavior associated with the development of complex morphologies in the ensemble. Other behaviors, *e.g.* cell adhesion, either do not generate forces, *i.e.* they only resist forces incoming from the environment or other cells or have a morphogenetic effect only when considering the mesenchyme. A major limitation of our study is that it does not include cell planar polarity. This polarity could have an effect on complexity and polarity. This possibility, however, should not affect our finding on how extracellular signaling can stabilize development and how complex morphologies can arise without such signaling.

Our results indicated that for complex morphologies to be developmentally stable, cell behaviors, specially cell contraction, should not be activated in the same way over large regions of the embryo. This heterogeneity can be achieved by gradients of gene expression, as in the gradient autonomous ensemble, or by extracellular signaling partitioning the embryo into different small regions of gene expression, as in the signaling ensemble. In either case cell behaviors can then be activated differently among contiguous regions and, thus, lead to more stable development. Cell signaling also allows for cells having specific patterns of gene expression (e.g. cell types) to be located in specific parts of a morphology. This is not possible in the autonomous and bio-mechanics-only ensemble.

The developmental mechanisms that can partition the embryo in such way are those that include the gene networks with extracellular signaling we found in the signaling-only ensemble. These are essentially the same networks found in other studies using the ensemble approach only with extracellular signaling (Salazar-Ciudad *et al*, 2000). In broad terms, these can be classified into Turing-like reaction-diffusion mechanisms (Turing, 1952) or hierarchic-signaling developmental mechanisms.

Our results on developmental stability suggest the existence of a strong constraint on the architecture of development. To ensure developmental stability, development should be structured in such a way that in no stage should there be large regions of cells regulating cell behaviors in the same way. Instead, the embryo should be undergoing constant partitioning so that, as it grows, the size of these regions remains small enough and most of the embryo is subdivided into such regions. This implies constant extracellular signaling as the embryo grows and deforms during morphogenesis. This simultaneity has been previously suggested for the animal development, although on different grounds (Salazar-Ciudad, 2003) and specially for vertebrates (Salazar-Ciudad *et al*, 2010).

Our results are also consistent with the large, and mostly random, morphological variability of embryoids (Simunovic & Brivanlou, 2017). In those we observe how the simple activation of cell behaviors can lead to relatively complex morphologies. Embryoids lack the spatially confined patterns of gene expression observed *in vivo* and that in our study are required for developmental stability (Simunovic & Brivanlou, 2017). This, together with the inevitably more noisy *in vitro* environment, may explain the large developmental instability of embryoids’ morphology.

It may seem paradoxical that relatively complex morphologies can be attained by the spatially homogeneous activation of cell behaviors, most notably contraction, over the whole embryo as in the autonomous and behaviors-only ensemble. This result can be understood by considering that the mechanical interactions between cells are usually non-linear (Oster & Alberch, 1982; Forgács & Newman, 2005; Taber, 2014) and, thus, as in Turing-like reaction-diffusion systems, simple homogeneous developmental patterns may be unstable to small perturbations on the system variables, *e.g.* cell positions, and easily change to more stable but spatially non-homogeneous configurations, e.g an invagination.

In more concrete terms, the arising of complexity from homogeneous initial developmental patterns can be understood, for example, by considering that the contraction of all cells in an epithelium will lead to a homogeneous increase in its curvature. This increase, for the geometrical reasons described in the results (see Figure S16), would resolve into the formation of several different invaginations in an embryo. As explained in the results, the position of each such invagination is largely affected by noise. The size and shape of each invagination, instead, is affected by the degree of cell contraction: the larger the contraction, the larger the curvature and more and smaller invaginations form. As invaginations form, they affect each others’ shape through mechanical interference and partial fusion. As a result of such interactions and the effect of noise, embryo’s epithelia do not fold into a regular array of invaginations but, instead, into an intricate sea of partially fused invaginations (as in Figure S9). These embryos are complex precisely because it is hard to predict the position of one cell based on the position of its neighbors.

Ours is not the only report of complex morphologies developing from homogeneous fields of cells that mechanically interact without extracellular signaling. Certain aspects of organ morphology, such as gut folding (Savin *et al*, 2011; Thomason *et al*, 2012; Nerurkar *et al*, 2017) or brain cortical folding (Bayly *et al*, 2013), have experimentally been shown to develop from homogeneous fields of cells without extracellular signaling being involved in the process. In more general terms, it has been argued that the development of many morphologies can be explained as a simple consequence of the mechanical properties of cells and their extracellular matrix, even from homogeneous initial conditions (Newman & Comper, 1990).

The relevance of extracellular signal gradients for the robust specification of the different regions of gene expression in embryos, *i.e.* “patterning”, has received extensive attention (Hogeweg, 2000). To our knowledge, however, their role in decreasing developmental instability at the level of morphogenesis has not been suggested before.

The questions we aim to address in this study could not have been addressed before, at least directly, because, until recently, there were no computational models that would be general enough to be applied to the development of any animal. There are currently a number of models that could fall in this category (Rejniak, 2007; Smith *et al*, 2012; Kriete & Eils, 2014). Ours is just one that is 3D, includes the different mechanical properties of epithelia and mesenchymal cells and, most importantly, it includes all animal cell behavior. Most of these general models have not been applied to general questions in evolution and development. The models used to address general questions in evolution and development tend to include only one cell behavior, extracellular signaling (Salazar-Ciudad *et al*, 2000; Kaneko, 2007; ten Tusscher & Hogeweg, 2011; Marcon *et al*, 2016) and, thus, do not consider that cells move and how morphology as such develops.

One exception is an article by Hogeweg (Hogeweg, 2000) using a 2D Potts model with a boolean gene network, extracellular signaling and cell adhesion to show that morphogenesis can evolve as a side effect of natural selection for diversity of cell types. The other exception is the work of Nissen *et al.* (Nissen *et al*, 2018). This work uses a 3D model including cell division, cell polarization and adhesion in epithelial cells without cell-cell extracellular signaling or gene networks. The initial conditions are a disorganized ball of epithelial cells that, over development time, organizes itself into a folded epithelium. Development, thus, does not occur, as in our model and in biology, through the activation of cell behaviors but is more like a phase separation event whose intermediate stages bear no direct resemblance to developmental stages.

There are several caveats related to our results. First, one may argue that complex animals are not just randomly folded epithelia and that, then, many of the morphologies we classify as complex do not necessarily resemble animal embryos. The way we measure complexity classifies seemingly random morphologies as more complex that morphologies that, somehow, seem more animal-like (see Figure S9). In fact, however, many existing measures of complexity, although not directly applicable to 3D morphology, understand that what seems random may indeed be quite complex (Kolmogorov, 1998; Wolfram, 2002). For us to do otherwise we would have to propose an unequivocal way to differentiate biological from non-biological complexity, a daunting task in itself. Anyway, both seemingly random and animal-like morphologies are classified as being rather complex according to our measures. In addition, our ensembles only consider small numbers of genes and cells and, thus, cannot include morphologies that look as complex as those of most actual animals (except perhaps for some sponges and cnidaria). In that respect, the inferences we will present on metazoan evolution should be regarded as primarily applying to the early metazoan evolution where, presumably, animals were made of relatively small numbers of cells. Notice that, in addition, we cannot use a measure of complexity that takes into account gene expression in space because in the autonomous ensembles, genes are, by definition, expressed homogeneously.

In any case, the result that the uniform regulation cell behaviors over large fields of cells leads to developmentally unstable morphologies does not depend on the way we measure complexity. The same applies to the result that the partitioning of the embryo into different regions of gene expression decreases developmental instability

Second, real developmental mechanisms are not random but the result of millions of years of evolution. It is precisely because we explored these mechanisms randomly, however, that we can claim that the requirements we found may be general: they should be fulfilled by the bulk of developmental mechanisms able to produce complex robust morphologies. One may argue that real developmental mechanisms are the result of a specific historical trajectory in the space of possible developmental mechanisms and that, then, a random sampling of such space, as in our ensembles, is not necessarily informative. However, our ensemble approach informs us about which of the requirements for complex robust morphologies are most commonly met in the space of possible developmental mechanisms and, thus, most likely to arise by mutation (Salazar-Ciudad *et al*, 2001). In other words, knowing which is the mutationally easiest way to lead to complex robust morphologies is informative about the possible path of evolution, at least as a null model on what to expect in the absence of other evolutionary forces.

Third, for computational reasons, we only simulated developmental mechanisms with up to 10 genes and stop simulations at embryos of 5000 cells. We think, however, that our conclusions would hold for ensembles considering more genes and cells: Large fields of cells activating the same cell behaviors would still tend to be unstable and partitioning such fields into smaller sub-fields would still tend to increase their developmental instability.

Fourth, one may wonder why to take an ensemble approach when one could just simulate evolution *in silico*, as we and others have done before for simpler models (Hogeweg, 2000; Salazar-Ciudad, Newman and Solé, 2001; Salazar-Ciudad and Marín-Riera, 2013; Vroomans, Hogeweg and ten Tusscher, 2016). We think that *in silico* evolution and the ensemble approach provide similar but complementary views on what is possible. The ensemble approach, however, is a computationally cheaper way to explore the space of possible developmental mechanisms (Kauffman, 1993). This is because in evolutionary simulations, most of the computational time is spent in simulating individuals that are closely related to each other (*i.e.* closer in the parameter space since they are relatives) while in the ensemble approach completely random individuals are simulated and then the space of possible developmental mechanisms is more evenly sampled. This does not preclude the future exploration of the more direct evolutionary approach but given the complexity of EmbryoMaker and its computational costs we found the ensemble approach more feasible.

Our results give support to existing theories on the evolution of early metazoa. According to Newman and Müller (Newman and Comper, 1990; Newman and Müller, 2000; Newman, Forgacs and Müller, 2006) early metazoans had relatively complex but very unstable morphologies. These authors argue that the behaviors and mechanical properties of animal cells allow for a relatively large repertoire of relatively complex morphologies. Strikingly this is what we found in the autonomous and cell behaviors-only ensembles: complex unstable morphologies arising by the activation of cell behaviors and mechanical interactions without extracellular signaling or even without complex gene regulatory networks. In addition, these authors argue that, later on, these complex and unstable metazoans evolved stable morphologies through the recruitment of complex gene networks in development. Although these authors do not provide much detail on how this happens, our results are consistent with the view that the early function of developmental gene networks and extracellular signaling may have been in stabilizing development rather than in building complex morphology *per se*.

The above argument by Newman and Müller concerns early metazoan evolution. In current metazoa, gene networks and extracellular signaling are pervasively important in the construction of morphology (Gilbert and Barresi, 2016). In addition, current complex metazoan morphology consist in something more than folded epithelia. These two facts suggest that, beyond earliest metazoan evolution, the role of gene networks and extracellular signaling would not be restricted to making complex morphologies stable but also to further increasing possible morphological complexity. This could be achieved by recombining existing developmental mechanisms in different stages and body parts. In other words, the use of gene networks and extracellular signaling allow a finer partitioning of the embryo, in each different stage, into territories and activate different developmental mechanisms in each of them. Although the basic morphologies are still those possible from the behaviors and mechanical properties of cells (such as rods, invaginations, cavities, etc.) (Newman and Comper, 1990; Newman and Müller, 2000) these are recombined through the different parts of the embryo to construct complex and slightly modular anatomies observed in many current metazoa. This possibility has not been simulated in this work because it requires computational resources still beyond our reach.

## Methods

A full description of EmbryoMaker can be found in its original publication (Marin-Riera *et al*, 2015). The cell behaviors considered in this article are: apoptosis, cell contraction (which can be asymmetric between the apical and basal side of epithelial cells and includes also cell expansion) and cell division (and accompanying cell growth) and extracellular matrix (ECM) secretion. In addition to cell behaviors, there are also a number of node mechanical properties such as their size, plasticity (the plastic reduction of cell’s size due to external pressure), cell adhesion affinities, resistance to compression and, for epithelial cells, resistance to epithelial bending. See SI for a more detailed description of the properties considered in this article.

### Building random developmental mechanisms

Random developmental mechanisms were built in four different ways. The set of morphologies that arise from each such ways we call an ensemble. In here we describe the basics of these ensembles.

All simulations started from the same simple initial developmental pattern: A flat hexagonal sheet of 126 epithelial cells and an underlying layer of 126 mesenchymal cells (see supplementary figure S2). Although EmbryoMaker allows for each cell to be made of several nodes, in this work, for simplicity, we consider that each epithelial cell was represented by a single cylindrical node and each mesenchymal cell by a spherical node. The initial values of the mechanical properties were the same in all cells in the initial developmental pattern. These initial values were such that, if unmodified over time, no changes in morphology will occur (see supplementary methods). Although the simulations included mesenchymal cells and ECM nodes, we did not consider them when measuring complexity and developmental instability. This is because the mesenchymal cells in the initial condition are not surrounded by an embryo-wide ectodermal epithelium and then can randomly spread over free space.

Each developmental mechanism was built by making randomly chosen genes to regulate randomly chosen genes, mechanical properties and cell behaviors (see supplementary figure S1). Developmental mechanisms were randomly built but they were not random over time: once a developmental mechanism is built it does not change over time. The gene network of each developmental mechanism represents only genetically-encoded potential regulations. Which gene products will interact in practice depends on which of the potentially interacting gene products will be present in each cell at the same time. The latter is not specified by the simulated developmental mechanism but arises from its dynamics over time.

When building the gene network of a developmental mechanism each gene had a 50% chance of being either an extracellularly diffusible gene product (in here we call these growth factors) or an intracellular gene product. The latter had a 0.25 probability to localize at the apical side of the epithelial cells (0.25 probability for the basal side) and a 0.5 probability to localize in both. We chose that gene 1 always directly activates a gene that can diffuse extracellularly. We specified that gene 1 always directly activates a gene that can diffuse extracellularly. We built gene networks of 10 genes by randomly wiring genes, each gene having a 0.2 probability of being a regulator of another gene in the network. Each regulation was, with equal chance, either positive or negative (transcriptional activation and repression) with a random regulative strength between 0 and *t*_*max*_ with uniform distribution. Thus, every gene had, on average, two positive and two negative connections (two efferent and two afferent). See supplementary material (SI 2.7.1) for a description of how is *t*_*max*_ chosen. We consider only transcriptional regulation between gene products, although EmbryoMaker can implement regulation at other levels and molecules other than gene products.

When building a developmental mechanism, each gene is given a chance to regulate a mechanical property or a cell behavior (with each gene having a 0.5 probability of doing so). The value of such regulation was random with an uniform or logarithmic distribution depending on the mechanical property and cell behaviors (see SI). Each gene product had a randomly chosen degradation and diffusion rate.

In addition, all cells had a default activation of cell division and cell differentiation that was constant over time (in addition of the regulation they could receive from a developmental mechanism). This reflects the fact that cell divisions take place in basically every developing embryo. Cell differentiation causes cell behaviors to slow down over developmental time. Including this cell differentiation is motivated by the widespread slowing down of growth during embryonic development.

All simulations were numerically integrated using the order 4 Runge-Kutta method with a dynamic step size. Simulations were run for a fixed number of iterations that is roughly equivalent to 3 physical days (see SI 1.7) unless largely aberrant morphologies were produced (*e.g.* consisting of broken epithelia). The number of iterations was chosen so that most embryos will finish development before this time. If largely aberrant morphologies were produced, *e.g.* broken epithelia, see SI 2.1.7 for a full description, these were considered inviable and discarded.

### The ensembles

We ran 100,000 random developmental mechanisms, we call *broad ensemble* to this set of random developmental mechanisms. This ensemble exhibited very few morphologies differing from the initial developmental pattern, and, thus, we devised a different way to build ensembles (see below). We found that nearly all random developmental mechanisms were unable to change gene expression patterns over space. Most genes were expressed only in the most central cell, in its immediate neighbor cells or not at all. As a result of this, most cells did not activate any cell behaviors nor did they change any mechanical property over time and space and, thus, there were no changes in cell positions and, thus, no morphogenesis.

To better identify random developmental mechanisms able to produce pattern transformations we built a simpler ensemble, the signaling-only *ensemble*, in which cells were not allowed to move, grow or divide. In this ensemble we identified which developmental mechanisms lead to changes in gene expression over space in a temporally stable fashion, as in a previous publication (Salazar-Ciudad *et al*, 2000). We then used the developmental mechanisms identified in such way to construct another ensemble, the signaling *ensemble,* by making some of the genes in each such mechanisms to regulate some randomly chosen node properties or cell behaviors. As a result, cells could move and morphogenesis occur, *i.e.* specific developmental patterns arose. The signaling ensemble is, thus, just a subset of the broad ensemble in which genes tend to be expressed beyond the central cell. In this ensemble, in addition, one gene (gene 1) was expressed in a gradient across the initial developmental pattern (figure 2).

We also constructed three additional ensembles that included the same developmental mechanisms (*i.e.* the same gene network, cell behaviors and node mechanical properties) than in the signaling ensemble but with different gene expression in the initial developmental pattern and no extracellular signaling.

*Autonomous ensemble* in this ensemble there is no extracellular signaling and one gene is homogeneously expressed through all cells in the initial developmental pattern while the rest are not expressed (see figure 2). Gene expression, thus, does not change in space but can change over time as a result of the model dynamics. Even if gene expression is homogeneous the bio-mechanical interactions between cells can lead to morphogenesis (a symmetry break is induced by noise and boundary conditions).

#### Autonomous bio-mechanics-only ensemble

This ensemble is like the autonomous ensemble but without a regulatory gene network. Thus, gene expression does not change in space nor in time but, as in the previous ensemble, morphogenesis can occur. Developmental mechanisms include only three genes, one is expressed in the apical part of cells, one in the basal side and one in both. These genes activate the cell behaviors that were regulated by the original developmental mechanism in the signaling ensemble and with the same intensity (see SI 2.5).

#### Gradient autonomous ensemble

This ensemble is just like the autonomous ensemble but only gene 1 is initially expressed and it is expressed in a gradient over the epithelium (see figure 2).

#### Complexity Measures

Our measures of complexity are related to predictability, *i.e.*, how likely it is to predict the position of an epithelial cell knowing the position of its neighbors cells. In a flat epithelium, for example, one can easily predict the position of a cell from the position of its closest neighbors since it would have the same position in the z axis. In a highly folded epithelium, this would be highly difficult, unless the epithelium happens to fold regularly, for example following a sinusoid wave. Thus, very complex morphologies are folded irregularly. See figure S4 to get an intuitive idea of each complexity measure and to see the complexity of several example morphologies. We use two different measurements of complexity:

#### Angle-distance variance (AV)

This measure is based on the variation of angles between epithelial cells. The angle between two cells is calculated as the angle between the apical-basal vectors of cell *i* and the apical-apical vector between cell *i* and *j* (see figure S6A).

To calculate AV, we first measure the angles formed between cell *i* and all other epithelial cells in a morphology. Then each cell is classified into one of seven categories based on its distance to cell *i* (see figure S6B). Each category falls into a specific distance interval, defined as follows:

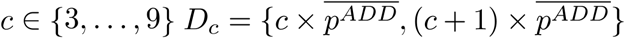

Where is the distance interval in which cell has to fall in to be included in the category *c*. *c* defines the maximal and the minimal distance for each interval. 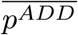 is the average *p*^*ADD*^ of all epithelial cells in the embryo. We start with *c* = 3 to preclude noise from affecting the measurement. We go up to *c* = 9, to take into account the macro-structure of the embryo.

Now we calculate the variance of the angles between cell *i* and the cells *j* that belong to a specific category *c*. We calculate the variance of each of the categories and add them together. We repeat these steps for all. The final angle variation complexity (AV) will be:

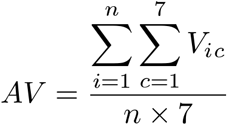

Where *i* is each of the epithelial cells, *n* is the total number of epithelial cells in the embryo, *c* is each of the categories intervals and *V*_*ic*_ is the angle variation for cell *i* in the category *c*. Notice that with this measurement a perfect sphere will have zero complexity.

#### Orientation Patch Count

(OPC) is based on the number of differently oriented slope patches an epithelium has. This measure is a fully 3D version of a measure of tooth complexity that has been found to correlate with diet (Evans *et al*, 2007).

For this method we first assign each epithelial cell to one of eight categories. Each category corresponds to one octant (one of the eight divisions of a Euclidean 3D coordinate system defined by the signs of the coordinates, see figure S7C). To determine in which octant the basal node is, we simply check the sign of each of the dimensions of the vector from the apical to the basal node of each epithelial cell.

Each cell is then further classified as belonging to a specific patch. A patch is a set of cells belonging to the same orientation category (of the 8 possible ones) and globally connected to each other. This means that one can go from any cell in a patch to any other cell in the patch through a sequence of contiguous cells belonging to same orientation category (see Figure S7B). By contiguous cells we mean cells that are in contact. Only patches with more than 3 cells were considered. Finally we simply count the number of patches in a morphology, which will give us the OPC value.

#### Developmental instability

We define developmental instability as the morphological distance between the morphologies resulting from simulating the same developmental mechanisms but with noise affecting how cells move. As with complexity measures we only take into account epithelial cells. We use two different complementary measures of morphological distance, the euclidean morphological distance and the orientation morphological distance (see SI 3.0).

#### Euclidean Minimal Distance (EMD)

This measure allows to compare morphologies made of different numbers of cells and without having to arbitrarily pre-select some landmarks of special morphological features (Salazar-Ciudad & Marín-Riera, 2013). This is very convenient for our study since embryos can be made or different numbers of cells. EMD is the mean distance from one node in a morphology to the closest node in another morphology. In other words, for each node in a morphology the distance to the closest node in the other morphology is calculated. Then the process is repeated for each node in the other morphology. All these distances are then averaged in respect to the number of nodes in one morphology and the other. In other words, between an arbitrary morphology 1 and an arbitrary morphology 2:

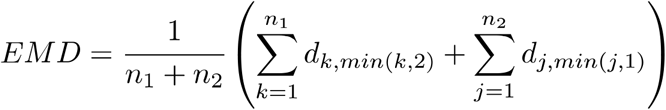

where *n*_*1*_ and *n*_*2*_ is the number of nodes in morphology 1 and 2 respectively, *d*_*k,min(k,2)*_ is the distance between node k in morphology 1 and its closest node in morphology 2, *d*_*j,min(j,2)*_ is the distance between node j in morphology 2 and its closest node in morphology 1. Note that that one node in morphology 1 is the closest node to another node in morphology 2 does not imply that this latter node is the closest to the former (these minimal distance relationships are not symmetric).

## Acknowledgements

We thank Miguel Brun-Usan, Lisandro Milocco, Renske Vroomans, Jukka Jernvall, for useful comments. This research was funded by the Finnish Academy to IS-C and CSC.

## References

1. Bonner JT (2004) Perspective: The size-complexity rule. Evolution 58(9):1883–1890.

2. Williams GC (1996) Adaptation and natural selection : a critique of some current evolutionary thought (Princeton University Press).

3. McCoy JW (1977) Complexity in organic evolution. J Theor Biol 68(3):457–8.

4. Hinegardner R, Engelberg J (1983) Biological complexity. J Theor Biol 104(1):7–20.

5. Gould SJ (2002) The structure of evolutionary theory (Harvard University Press).

6. Arendt D (2008) The evolution of cell types in animals: emerging principles from molecular studies. Nat Rev Genet 9(11):868–882.

7. Canning EU, Okamura B (2004) Biodiversity and evolution of the Myxozoa. Adv Parasitol 56:43–131.

8. Arthur W (2011) Evolution : a developmental approach (Wiley-Blackwell)

9. Fisher DC (1986) Progress in Organismal Design. Patterns and Processes in the History of Life, eds Raup DM, Jablonski D. (Springer, Heidelberg), pp 99–117.

10. Ruse M (2009) Monad to man : the concept of progress in evolutionary biology (Harvard University Press).

11. McShea DW (1996) Perspective metazoan complexity and evolution: is there a trend? Evolution (N Y) 50(2):477–492.

12. Fleming L, Mcshea DW (2013) Drosophila mutants suggest a strong drive toward complexity in evolution. Evol Dev. 15(1):53–62.

13. Salazar-Ciudad I (2003) Mechanisms of pattern formation in development and evolution. Development 130(10):2027–2037.

14. Davies JA (2013) Mechanisms of morphogenesis (Elsevier Academic Press).

15. Newman SA, Bhat R (2009) Dynamical patterning modules: a “pattern language” for development and evolution of multicellular form. Int J Dev Biol 53(5–6):693–705.

16. Gilbert SF, Barresi MJF (2016) Developmental biology (Oxford University Press)

17. Barkoulas M, van Zon JS, Milloz J, van Oudenaarden A, Félix MA (2013) Robustness and Epistasis in the C. elegans Vulval Signaling Network Revealed by Pathway Dosage Modulation. Dev Cell. 24(1):64–75.

18. Oliveira MM, Shingleton AW, Mirth CK (2014) Coordination of Wing and Whole-Body Development at Developmental Milestones Ensures Robustness against Environmental and Physiological Perturbations. PLoS GeneAt. 10(6) e1004408.

19. Marin-Riera M, Brun-Usan M, Zimm R, Välikangas T, Salazar-Ciudad I (2015) Computational modeling of development by epithelia, mesenchyme and their interactions: a unified model. Bioinformatics. 32(2):219–25.

20. Saunders WB, Work DM, Nikolaeva SV (1999) Evolution of Complexity in Paleozoic Ammonoid Sutures. Science. 286,760–763.

21. Evans AR, Wilson GP, Fortelius M, Jernvall J (2007) High-level similarity of dentitions in carnivorans and rodents. Nature 445(7123):78–81.

22. Arjan J, et al. (2003) Evolution and detection of genetic robustness. Evolution. 57(579):1959–1972.

23. Salazar-Ciudad I (2007) On the origins of morphological variation, canalization, robustness, and evolvability. Integr Comp Biol 47(3):390–400.

24. Waddington ch (1942) Canalization of development and the inheritance of acquired characters. Nature 150(3811):563–565.

25. Shapiro BL (1971) Developmental stability and instability. J Dent Res 50(6):1505–6.

26. Klingenberg CP (2002) Morphometrics and the role of the phenotype in studies of the evolution of developmental mechanisms. Gene 287(1–2):3–10.

27. Salazar-Ciudad I, Garcia-Fernández J, Solé R V (2000) Gene networks capable of pattern formation: from induction to reaction-diffusion. J Theor Biol 205(4):587–603.

28. Salazar-Ciudad I, Marín-Riera M (2013) Adaptive dynamics under development-based genotype-phenotype maps. Nature 497(7449):361–4.

29. Martin AC, Goldstein B (2014) Apical constriction: themes and variations on a cellular mechanism driving morphogenesis. Development 141(10):1987–98.

30. Bunn JM, et al. (2011) Comparing Dirichlet normal surface energy of tooth crowns, a new technique of molar shape quantification for dietary inference, with previous methods in isolation and in combination. Am J Phys Anthropol 145(2):247–261.

31. Striedter GF, Srinivasan S, Monuki ES (2015) Cortical Folding: When, Where, How, and Why? Annu Rev Neurosci 38(1):291–307.

32. Bard JBL (1992) Morphogenesis : the cellular and molecular processes of developmental anatomy (Cambridge University Press).

33. Turing A (1952) The chemical basis of morphogenesis. Philos Trans R Soc Lond B Biol Sci 237(641):37–72.

34. Salazar-Ciudad I, et al. (2010) Morphological evolution and embryonic developmental diversity in metazoa. Development 137(4):531–539.

35. Simunovic M, Brivanlou AH (2017) Embryoids, organoids and gastruloids: new approaches to understanding embryogenesis. Development 144(6):976–985.

36. Oster G, Alberch P (1982) Evolution and Bifurcation of Developmental Programs. Evolution 36(3):444–459.

37. Forgacs G, Newman SA. (2005) Biological physics of the developing embryo (Cambridge University Press).

38. Taber LA (2014) Morphomechanics: transforming tubes into organs. Curr Opin Genet Dev 27:7–13.

39. Savin T, et al. (2011) On the growth and form of the gut. Nature 476(7358):57–62.

40. Thomason RT, Bader DM, Winters NI (2012) Comprehensive timeline of mesodermal development in the quail small intestine. Dev Dyn 241(11):1678–94.

41. Nerurkar NL, Mahadevan L, Tabin CJ (2017) BMP signaling controls buckling forces to modulate looping morphogenesis of the gut. Proc Natl Acad Sci U S A 114(9):2277–2282.

42. Bayly P V., Okamoto RJ, Xu G, Shi Y, Taber LA (2013) A cortical folding model incorporating stress-dependent growth explains gyral wavelengths and stress patterns in the developing brain. Phys Biol. 10(1):016005.

43. Newman SA, Comper WD (1990) “Generic” physical mechanisms of morphogenesis and pattern formation. Development 110(1):1–18.

44. Hogeweg P (2000) Evolving Mechanisms of Morphogenesis: on the Interplay between Differential Adhesion and Cell Differentiation. J Theor Biol 203(4):317–333.

45. Rejniak KA (2007) Modelling the Development of Complex Tissues Using Individual Viscoelastic Cells. Single-Cell-Based Models in Biology and Medicine (Birkhäuser Basel, Basel), pp 301–323.

46. Smith AM, Baker RE, Kay D, Maini PK (2012) Incorporating chemical signalling factors into cell-based models of growing epithelial tissues. J Math Biol 65(3):441–463.

47. Kriete A, Eils R (2014) Computational systems biology (Elsevier/AP).

48. Kaneko K (2007) Evolution of Robustness to Noise and Mutation in Gene Expression Dynamics. PLoS One 2(5):e434.

49. ten Tusscher KH, Hogeweg P (2011) Evolution of Networks for Body Plan Patterning; Interplay of Modularity, Robustness and Evolvability. PLoS Comput Biol 7(10):e1002208.

50. Marcon L, Diego X, Sharpe J, Müller P (2016) High-throughput mathematical analysis identifies Turing networks for patterning with equally diffusing signals. Elife 5:e14022.

51. Nissen SB, Rønhild S, Trusina A, Sneppen K (2018) Theoretical tool bridging cell polarities with development of robust morphologies. Elife 7:e38407.

52. Kolmogorov AN (1998) On tables of random numbers. Theor Comput Sci 207(2):387–395.

53. Wolfram S (2002) A new kind of science (Wolfram Media).

54. Salazar-Ciudad I, Newman SA, Solé R V (2001) Phenotypic and dynamical transitions in model genetic networks. I. Emergence of patterns and genotype-phenotype relationships. Evol Dev 3(2):84–94.

55. Vroomans RMA, Hogeweg P, ten Tusscher KHWJ (2016) In silico evo-devo: reconstructing stages in the evolution of animal segmentation. Evodevo 7(1):14.

56. Kauffman SA (1993) The origins of order : self-organization and selection in evolution (Oxford University Press).

57. Newman SA, Müller GB (2000) Epigenetic mechanisms of character origination. J Exp Zool 288(4):304–317.

58. Newman SA, Forgacs G, Müller GB (2006) Before programs: The physical origination of multicellular forms. Int J Dev Biol 50(2–3):289–299.

