## Supplementary for "Cell signalling stabilizes morphogenesis against noise"


1. The EmbryoMaker modeling framework
  1.1 Introduction to EmbryoMaker
  1.2 Bio-mechanics
  1.3 Gene expression and gene networks
  1.4. Cell-cell signaling
  1.5. Regulation of node properties
    1.5.1 Regulation of node properties
    1.5.2 The regulation of node radii
  1.6. Cell behaviors
1.7 Model time
1.8 Summary of model parameters

2. The ensemble approach
  2.1 The broad ensemble
    2.1.1 Construction of gene networks
    2.1.2 Diffusion coefficient
    2.1.3 Degradation rate
    2.1.4 Gene regulation of node mechanical properties and behaviors
    2.1.5. Initial conditions
    2.1.6 Simulation of each developmental mechanism
    2.1.7. End of development simulation.
    2.1.8. Criteria for inviable embryos
  2.2. Signaling-only ensemble
    2.2.1 Construction of gene networks
    2.2.2 Diffusion coefficient
    2.2.3 Degradation rate: Just as in the broad ensemble, section 2.1.3
    2.2.4 Gene regulation of node mechanical properties and behaviors
    2.2.5. Initial conditions
    2.2.6 End of development simulation
    2.2.7 Classifications of the resulting patterns
  2.3 Signaling ensemble
    2.3.1 Gene regulation of node mechanical properties and behaviors
    2.3.2 Initial conditions
    2.3.4 Simulation of each developmental mechanism
    2.3.5 End of development simulation
  2.4 Autonomous ensemble
  2.5 Autonomous bio-mechanics-only ensemble
  2.6 Gradient autonomous ensemble
  2.7 Ranges of the model parameters
    2.7.1 Maximal transcription regulation strength per gene, $t_{max}$
    2.7.2 Diffusion coefficient
    2.7.3 Degradation rate:
    2.7.4 Regulation of node properties, $e_{k,min}$ and $e_{k,max}$
      2.7.4.1 Components of $p^{EQD}$: $p^{COD}$, $p^{GRD}$, $p^{PLD}$ and $p^{VOD}$
      2.7.4.3. $p^{ADD}$, adhesion radius
      2.7.4.4. $p^{YOU}$: intracellular elasticity
      2.7.4.5. $p^{REC}$, cell compressibility, $p^{ADH}$, intercellular adhesion, $p^{HOO}$, apico-basal elasticity and the epithelial torsion properties $p^{ERP}$ and $p^{EST}$
      2.7.4.6. $p^{EQS}$, Apico-basal equilibrium distance
      2.7.4.7. $p^{MOV}$, Filopodial instability
      2.7.4.8. $p^{DMO}$, Filopodia extensibility

2.7.4.9. $p^{DIF}$, cell differentiation
2.7.5 Regulation of cell behaviors, $c_{k,min}$ and $c_{k,max}$
2.7.5.1. Cell division
2.7.5.2. Apoptosis
2.7.5.3 Epithelium-to-mesenchyme transition (EMT):
2.7.5.4. ECM secretion:

2.8. Node mechanical property values in the initial conditions

3. Orientation Morphological Distance (OMD)
3.1 *Orientation Morphological Distance (OMD)*
3.2 Flattening of the epithelium

4. Other analysis.
4.1 Disparity
4.2 Developmental stability
4.3 number of cells contracting in the same way
5. References

#### 1. The EmbryoMaker modeling framework:

##### 1.1 Introduction to EmbryoMaker:

The development model we use in this article is EmbryoMaker (Marin-Riera *et al*, 2015). Except for a small detail, explained in section 1.5.1, everything is done with EmbryoMaker version 1.0, http://www.biocenter.helsinki.fi/salazar/software.html. The following section describes the basics of EmbryoMaker and how we use it. For a more detailed description see (Marin-Riera *et al*, 2015).

EmbryoMaker is a programming framework for animal development. It is specially designed to easily construct mathematical models of pattern formation and morphogenesis in animal development. It is not a model in the sense that it does not specify which genes or which cells should interact during development to produce a specific morphology. EmbryoMaker specifies, instead, some generic equations about how gene products and cells interact and some rules about how to implement basic cell behaviors (such as cell division, apoptosis, etc…). The actual number of gene products and the intensity of their interactions is not specified by EmbryoMaker but by each specific model build using it. Any model build with EmbryoMaker is also a specification of how some of these gene products regulate specific cell mechanical properties and cell behaviors. Thus, while EmbryoMaker provides the generic equations, each system-specific model provides the values of the parameters in such equations and the number of such equations (since there are several equations per gene product, cell mechanical properties and behaviors). In this way mathematical models specific of spiral cleavage (Brun-Usan *et al*, 2017) and early tooth development (Marin-Riera *et al.*, 2018) have been constructed with EmbryoMaker.

EmbryoMaker includes a set of variables and a set of parameters. How these variables change over time from their values in the initial conditions is what one wants to understand or predict from each specific model. How these variables change depends on: their initial values, the set of equations describing gene product interactions, the set of equations describing cell interactions, on cell behaviors and on the values of the parameters in these equations. The values of the parameters in these equations and values of the variables in the initial conditions are specified by the user to make models for specific developmental systems. The actual number of equations in each model is also decided by the user since the number of gene products, types of cell mechanical interaction and cell behaviors used in a specific model is also decided by the user (from a large pre-specified set).

Variables specify for example, the position in 3D of a cell or cell part, the concentration of a gene product in a cell or cell part and also some mechanical properties of cells, like their adhesivity, that may change over time as a result of changes in gene product concentrations. Most of these variables are associated with a cell or cell part (*e.g.* the x coordinate of a given cell, how much of a specific gene product this cell expresses, or its unspecific adhesivity). Cells are made of parts that we call nodes, the position of a cell's nodes in space defines its shape. Each node has a set of mechanical properties and levels of expression for each gene considered in each specific model. All these are variables of the model and as such can change over model's simulation time. In here we denote each node mechanical property with a p and superscript. Thus, $p^{EQD}{}_i$ is, for example, the node mechanical property *EQD* of node *i*. Some mechanical properties apply only to whole cells and then we denote them in the same way but using a capital p, *P*. Notice that he number of variables, but not the number of parameters, would usually change during a simulation since each cell has a number of variables (e.g. its position in 3D) and the number of cells can change as a result of cell division or cell death. The number of variables and their initial values are specified in the initial conditions in each specific model. Each initial condition is simply a set of cells with a specific distribution in 3D space and a specification of the gene expression and mechanical properties of each cell. Each initial condition is simply an embryo, organ or multicellular aggregate in a specific stage. Even if EmbryoMaker can simulate any kind of multicellular system, in here we use the term embryo for each system simulated by EmbryoMaker.

Parameters do not change during a simulation, variables do, and they are supposed to be genetically encoded. Model parameters specify for example, how strongly a gene product regulates the transcription of another one, how strongly a gene product promotes some specific cell behavior or the diffusion coefficient of a specific gene product. Note that how much a transcriptional factor A is transcribed in a given cell and moment as a result of being regulated by transcriptional factor B is determined by the dynamics of the model (it is a variable) but the binding affinity of transcriptional factor B for the promoter of A is a parameter of the model (*i.e.* it is genetically encoded) and, thus, is given from outside (*i.e.* specified by the user when implementing a specific model). The set of cross-regulations between gene products we call a model's gene network. When we consider a gene network and how genes in it regulate cell behaviors or mechanical properties, we talk of developmental mechanisms, since for pattern formation to occur there has to be changes in cell behaviors or mechanical properties, see (Salazar-Ciudad, 2003). Notice that the sets of gene products and gene interactions involved in development may change over developmental stages but that the gene network, as we define it in here, will not. The gene network includes all possible genetically encoded interactions between gene products. As a result of each model dynamics, these possible interactions may occur or not depending on whether the involved genes happen to be expressed in the same cells at the same time (this will depend on how cells have moved, which signaling from other cells they received over time, etc.).

##### 1.2 Bio-mechanics:

EmbryoMaker can simulate mesenchymal and epithelial cells in addition to extracellular matrix (ECM) and the mechanical interactions between them all. Mesenchymal cells are made of spherical bodies, that we call nodes, whereas epithelial cells are made of cylindrical bodies consisting of two nodes (one basal and one apical bound by an elastic link) (Figure 1). Although EmbryoMaker can simulate cells made of any number of nodes, in the present article we consider only the case in which each mesenchymal cell is made of a single spherical node while each epithelial cell is made of a single cylinder, since this greatly increases the speed of the simulations. ECM is made of spherical nodes that do not belong to any cell. The movement of nodes follows an over-damped Langevin equation of motion:

$$\frac{\partial \vec{r}_i}{\partial t} = \sum_{j=1}^{j=n_v} f_{Aij} \ \hat{u}_{ij} \qquad (1)$$

where $r_i$ is the position of node $i$ in three-dimensional space, $n_v$ is the number of nodes that are close enough to node $i$ to mechanically interact with it, $t$ is time, $f_{Aij}$ is the modulus of the force acting between node $i$ and $j$ and $u_{ij}$ is the unit vector connecting node $i$ and node $j$ (see Figure S3). The modulus and sign of the force is dependent on the distance between the two nodes:

$$\begin{cases} f_{Aij}=\left(p_i^{REC}+p_j^{REC}\right)\left(d_{ij}-\left(p_i^{EQD}+p_j^{EQD}\right)\right) \ \ if \ \ d_{ij}<\left(p_i^{EQD}+p_j^{EQD}\right) \\ f_{Aij}=k_{ij}^{ADH}\left(d_{ij}-\left(p_i^{EQD}+p_j^{EQD}\right)\right) \ \ if \ \ \left(p_i^{EQD}+p_j^{EQD}\right)\leq d_{ij}\leq\left(p_i^{ADD}+p_j^{ADD}\right) \\ f_{Aij}=0 \ \ if \ \ \left(p_i^{ADD}+p_j^{ADD}\right)>d_{ij} \end{cases} \quad (2)$$

When the distance between nodes $i$ and $j$ ($d_{ij}$) is shorter than the sum of their radii at equilibrium (node property $p^{EQD}$), there is a repulsive force proportional to the sum of the node property $p^{REC}$ of each node (this coefficient determines their incompressibility). When this distance is longer than the equilibrium distance but shorter than the sum of the maximum radii of $i$ and $j$ (node property $p^{ADD}$), there is an attractive force between nodes i and j. This force is proportional to $k_{ij}^{ADH}$,

$$k_{ij}^{ADH} = g_{im}g_{jn}b_{mn} \qquad (3)$$

where $g_{im}$ is the amount of adhesion molecule $m$ expressed in node $i$ and $b_{mn}$ is the adhesive affinity between adhesion molecules $m$ and $n$. The set of binding affinities between each pair of adhesion molecules is contained in the $B$ matrix.

The direction of force vectors differ between mesenchymal-mesenchymal, epithelial-epithelial and the epithelial-mesenchymal node interactions, since vectors need to be normal to the contact interface between nodes and nodes have different shapes in epithelial cells and mesenchymal cells (see (Marin-Riera *et al*, 2015) for a detailed explanation).

The apical and basal nodes of epithelial cells are connected by an elastic spring that opposes any departure from an equilibrium distance between the apical and basal nodes of each cylinder (14). The force generated by the spring is calculated as follows,

$$\overrightarrow{f}_{sij} = k_{ij}^{HOO}(d_{ij} - p_{ij}^{EQS})\hat{s}_{ik} \qquad (4)$$

where $k_{ij}^{HOO} = p_i^{HOO} + p_j^{HOO}$ is the elastic coefficient of the spring (which is determined by the sum of the mechanical parameter $p^{HOO}$ in both nodes), $d_{ij}$ is the distance between node $i$ and $j$, $p_{ij}^{EQS}$ is the equilibrium length of the spring between node $i$ and $j$ and $s_{ik}$ is the unit vector connecting the two epithelial nodes.

Two additional force components are applied to epithelial cells in order for them to organise as one layered sheets. A radial force acts along the apical-basal axis of the cell and tends to restore displacements in that axis in respect to neighboring cells in the epithelium, whereas a rotational force acts tangential to surface of the epithelium and tends to orient the apical-basal axis of cells normal to the epithelial plane. These forces are calculated as follows,

$$\overrightarrow{f}_{ESTij} = k_i^{EST}\frac{\overrightarrow{m}_{ijkl}\cdot\overrightarrow{c}_{ij}}{|\overrightarrow{m}_{ijkl}|}\hat{m}_{ijkl} \qquad (5)$$

$$\overrightarrow{f}_{ERPij} = p_i^{ERP}\frac{\overrightarrow{s}_{ik}\cdot\overrightarrow{c}_{ij}}{|\overrightarrow{s}_{ik}|}\hat{c}_{ij} \qquad (6)$$

where $f_{ij}^{EST}$ is the radial bending force and $f_{ij}^{ERP}$ is the rotational bending force. We define $c_{ij}$ as the vector connecting neighboring node i and j, $s_{ik}$ and $s_{jl}$ as the vectors that connect each apical node to their basal counterparts and $m_{ijkl}$ as the sum of $s_{ik}$ and $s_{jl}$ which defines the vector normal to the apical or basal surface between i and j. The radial bending force always acts on the direction of $m_{ijkl}$, and is proportional to the deviation of the angle formed by $m_{ijkl}$ and $c_{ij}$ from 90º. $k_{ij}^{EST}$ is the sum of the mechanical parameter $p^{EST}$ of nodes *i* and *j* (see (14) for a detailed explanation). The rotational bending force is proportional to the deviation of the angle formed by $s_{ik}$ and $c_{ij}$ from 90º, but in this case the direction of the force is parallel to $c_{ij}$, thus promoting a tilting of the epithelial cylinder that reaches an equilibrium (that is the force modulus becomes 0) when the apical-basal axis of the epithelial cylinder is normal to the apical/basal cell surface. $k_{ij}^{ERP}$ is the sum of the mechanical parameter $p^{ERP}$ of nodes *i* and *j*. (see (14) for a detailed explanation).

In summary, thus, the forces acting on an epithelial node are:

$$\frac{\partial \vec{r}_i}{\partial t} = \vec{f}_{sik} + \sum_{j=1}^{n_d} \left( f_{Aij}\hat{u}_{ij} + \vec{f}_{ESTij} + \vec{f}_{ERPij} \right) \quad (7)$$

where *k* is the node in the same cylinder than *i* and the sum is made over all the neighboring nodes except for *k*.

In addition to the movement equation defined at (7) there is noise in node movements. At each time step, a proportion $M_{NOI}$ (a global model parameter) of the nodes are chosen at random and are tentatively moved in a random direction for a random distance between 0 and $p^{DMO}_i$, a mechanical property of each node. For each node the potential mechanical energy is calculated, by integrating the same force equation (1), in the new position. If the potential energy in the new position is smaller than in the old position the movement is accepted. If not, the movement is accepted with a probability proportional to the difference in potential energy between the new and old positions and inversely proportionally to a temperature parameter, model parameter $M_{TEM}$, plus a node property defining the node's propensity to movement ($p^{MOV}$). If the movement is not accepted the node is put back to its old position. This energy biased noise reflects the fact that noise can affect nodes' positions but it is unlikely to bring nodes into very energetically unfavorable positions (*e.g.* noise is very unlikely to bring a node from a cell inside another cell). This is a standard way to implement noise in many physical and biological systems ,such as in the Pott's model (Graner & Glazier, 1992). Except temperature and noise proportion, all the quantities defined so far are variables that are specified in the initial conditions and may change as a result of model dynamics.

##### 1.3 Gene expression and gene networks

EmbryoMaker considers gene products but also other kinds of molecules that are not transcribed. In this article, for simplicity, we consider only gene products and only transcriptional regulation. Each gene product has a set of properties associated with it (which we call genetic parameters). These include its intrinsic degradation rate ($\mu_i$) and how they affect transcription, node properties and cell behaviors. The rate of change in the concentration of gene *k* in node *i* over time is:

$$\frac{\partial g_{ik}}{\partial t} = \frac{\phi\left(\sum_{l=1}^{n_g} t_{lk} g_{il}\right)}{1 + \phi\left(\sum_{l=1}^{n_g} t_{lk} g_{il}\right)} - \mu_k g_{ik} \quad (8)$$

where $\mu_k$ is the intrinsic rate of degradation of molecule *k*, $g_{il}$ is the amount of transcriptional factor *l* in node *i* and each $t_{lk}$ term is the strength by which each specific transcriptional factor *k* activates (positive $t_{lk}$) or inhibits (negative $t_{lk}$) the transcription of gene *l*. The sum is done through all the regulatory molecules and by definition only transcriptional factors have $t_{lk}$ terms different from zero. $\phi$ is a function that is equal to 0 for values of x smaller than 0 and equals to x when x is greater than 0 ($\phi$(x)=0 if x<0 and $\phi$(x)=x if x>0). This function is used to ensure that there is not such a thing as negative transcription (although $t_{lk}$ can be negative and thus repress transcription). Each $t_{lk}$ is a model parameter, the set of all the possible $t_{lk}$ is what we call the T matrix, a $n_g$ x $n_g$ matrix where $n_g$ is the number of gene products in a model. Matrix $T$, defines a model's gene network.

Equation (8) represents the binding of several transcriptional factors to the promoter of gene $k$. This is a saturating process that, for simplicity, is represented by a Hill equation of order 1. This means that when there are few activator factors the rate of transcription increases with the amount of these factors. But when there are many of these factors the rate of transcription does not increase as much with the amount of activator factors since the binding sites are likely to be already occupied. Equation (8) also implies that the maximal rate of transcription of a gene is 1. The same (Salazar-Ciudad *et al*, 2000, 2001),) or similar(Reinitz & Sharp, 1995) equation has been used in previous models of gene networks in development.

##### 1.4. Cell-cell signaling:

In EmbryoMaker some gene products can be chosen to diffuse in the extracellular space between cells. EmbryoMaker considers sophisticated ligand-receptor dynamics but in this article we consider the most simple ideal case in which diffusible gene products could affect transcription directly (so we consider each signal transduction pathway to be transmitting signal in a lineal perfect way without amplification). Diffusion is implemented as transfers of molecules between nodes (including ECM nodes). This transport follows Fick's second law of diffusion:

$$\frac{\partial q}{\partial t} = -D \bigtriangledown^2 q \qquad (9)$$

Where $q$ is concentration of a molecule, $D$ is the diffusion coefficient of that molecule and $\bigtriangledown^2 q$ is the second derivative of the concentration in 3D space. We calculate transfers of matter between pairs of nodes. Since we only make calculations in the nodes, diffusion is essentially discrete (although non-uniformly) and this equation is roughly approximated by:

$$d_{ik} = D_k \sum_{j=1}^{n_v} \left( \frac{g_{ik} - g_{jk}}{d_{ij}} \right) \quad (10)$$

Where $g_{ik}$ is the amount of molecule $k$ in node $i$, $t$ is time, $D_k$ is the diffusion coefficient of molecule $k$, $n_v$ is the number of nodes within the maximum radius of diffusion from node $i$ and $d_{ij}$ is the distance between node $i$ and $j$. Both this distance and $n_v$ depend on how nodes are arranged in space. The maximum radius of diffusion is two times the maximal $p^{ADD}$. This latter choice ensures an optimal accuracy even if there are changes in the sizes of the nodes in the embryo over time (see (Marin-Riera *et al*, 2015) for details). The change in concentration of diffusible molecule $k$ in node $i$ over time is then:

$$\frac{\partial g_{ik}}{\partial t} = \frac{\phi\left(\sum_{l=1}^{n_g} t_{lk} g_{il}\right)}{1 + \phi\left(\sum_{l=1}^{n_g} t_{lk} g_{il}\right)} - \mu_k g_{ik} + d_{ik} \quad (11)$$

##### 1.5. Regulation of node properties

*1.5.1 Regulation of node properties:*

The most relevant node properties have already been described when describing mechanical forces. See section 1.5.3 for a list of those. Each node property value in a node can be modified by the amounts of specific regulatory molecules in a node. In EmbryoMaker the value of node property *l* at time *t* in node *i* is then:

$$p_i^l(t) = \phi\left(p_i^l(0) + (1 - p_i^{DIF})\sum_{k=1}^{n_g} e_{lk} g_{ik}\right) \quad (12)$$

Where $p_i^l(t)$ is the value of node property *l* in node *i* at time *t* and $p_i^l(0)$ is the value of that node property *l* in node *i* when the node was created (this is in the initial condition or when the node first arose through cell division). Fuction $\phi$, as in equation (8), ensures that node properties can become very small (or zero) but not negative. $p_i^{DIF}$ is the degree of differentiation in node *i* (differentiation slows down changes in nodes). The amount of change in node properties is then related to how much of the molecules regulating these properties there is in a node and how strongly they regulate them, as specified in each element *e*. Each element *e* is a model parameter. The set of all elements *e* in a model is the *E* matrix. In EmbryoMaker this regulation is supposed to be instantaneous compared with the rate at which nodes move or with the rate at which regulatory molecules are catalyzed. In this article, however, the rate of change of each node property per model time unit is not allowed to be larger than 0.5% of node property value at the initial conditions. This is the only difference between the original EmbryoMaker (Marin-Riera *et al*, 2015) and the version we use in here. A full description of all node properties can be found in the original model description (Marin-Riera *et al*, 2015). Section 2.6.4 and 2.6.5 also provide some short description.

*1.5.2 The regulation of node radii:*

The node property $p^{EQD}$ is the maximal distance, from a node's *i* center, at which another node has to be to experience a repulsion force from node *i*. At each time instant this node property is equal to the sum of four other node properties that correspond to four different cell processes.

$$p_i^{EQD} = p_i^{COD} + p_i^{GRD} + p_i^{PLD} + p_i^{VOD} \quad (13)$$

The first term is coming from node active contraction (due to myosin and related molecules), the second is coming from cell growth and apoptosis, the third from cell mechanical plasticity and the fourth from cylinder volume conservation. Having $p^{EQD}$ determined by four independent terms allows contraction, growth, plasticity and volume conservation to occur at the same time in a cell. For example it is important that parts of the cell can contract while the cell is growing (and that would not be possible if growth and contraction would act directly on $p^{EQD}$ since then growth would increase $p^{EQD}$ and contraction would decrease it). Cell contraction is realized when a gene product negatively regulates node property $p^{COD}$.

##### 1.6. Cell behaviors

EmbryoMaker includes a number of cell behaviors. As in animal cells, these can be regulated genetically. The C matrix quantitatively specifies how each gene product regulates each cell behavior, analogously to the E matrix for node properties. A specific cell variable exist for each cell behavior and a different equation exists for each cell behavior. .

All cell behaviors are implemented as simple logical operations on nodes. Cell division is implemented by placing a new cell in a random position close to plane of division of the cell (which is normal to the longest axis of the cell, see (Marin-Riera *et al*, 2015)). Cell contraction is implemented in the model by changes in the node property $p^{COD}$ as explained previously. As depicted in equation (2) and (3), cell adhesion is integrated in the mechanical part the model. Each node has a basal adhesivity plus the one given by the expression of adhesion molecules, which depends on the affinity of the adhesion molecules expressed in each node (represented in the B matrix, see (Marin-Riera *et al*, 2015)). This includes also the possibility to implement repulsion between cells (negative values in B matrix elements). Apoptosis is implemented by a gradual decrease in cell radius (through changes in $p^{GRD}$ ) until the cell disappears. Epithelial cells can also be induced to undergo an epithelial-mesenchymal transition. Both mesenchymal and epithelial cells can secrete ECM nodes.

##### 1.7 Model time:

There are two different kinds of time in the simulations using EmbryoMaker. The actual computation time that it took to simulate each developmental mechanism. Then there is the actual amount of development time simulated (e.g. 10 hours of embryonic development). This time is a continuous variable. The time units are hours but to preclude confusion between the two times we talk about model time units instead of hours. Different amounts of computational time are usually required to simulate the same amount of developmental time for different developmental mechanisms. The actual computational time depends on how many cells an embryo has over time, how many different cell behaviors, genes and gene interactions occur during its development. Computational and developmental time depend additionally on how the model is numerically integrated, which depend on some of the global parameters of the model. In this article, all simulations numerical integration used the order 4 Runge-Kutta method with a dynamic step size.

##### 1.8 Summary of model parameters:

EmbryoMaker is a model of models. By itself it does not specify much about any specific development other than the generic equations for gene product regulation, node biomechanics, cell behaviors and a set of possible node mechanical properties. The actual mathematical models of specific systems development are built by specifying a gene network and how the genes in it regulate specific node properties and cell behaviors. This is the same than specifying the size and values in the T,E,C and B model parameter matrices and the degradation rates and diffusion coefficients for each gene product (the M and D vectors). In fact, in here, we call each combination of specific T, E, C, B matrices and M and D vectors a developmental mechanism since it is not just a gene network but a specific link between it and bio-mechanics. Each element in these matrices and vectors is a parameter of the model. Note, however, that the model is very flexible and does not presuppose any number of genes, that is specific to each actual model. This implies that the number of parameters and the sizes of these matrices and vectors is not fixed by EmbryoMaker but depends on each specific model. Thus, if the network of a specific model with EmbryoMaker has $n_g$ genes the T matrix would be a $n_g$ x $n_g$ matrix, the E matrix would be a $n_g$ x $n_p$ matrix and the C matrix would be a $n_g$ x $n_c$ matrix, where $n_p$ is the number of node properties considered by EmbryoMaker and n $n_c$ the number of cell behaviors considered by EmbryoMaker. These last two numbers are fixed and specific of EmbryoMaker, although in practice many models would only make use of some small specific subset of those node properties and cell behaviors (e.g. the tooth model we publish with EmbryoMaker considers only cell division and adhesion).

#### 2. The ensemble approach

In this section we detail how the different ensembles were built. This is, how random gene networks were built, and how gene products in those were chosen to regulate node properties and cell behaviors. We also describe the initial conditions used, how was each simulation run and which criteria were used to stop development in each simulation.

##### 2.1 The broad ensemble:

*2.1.1 Construction of gene networks*

We considered gene networks of 10 gene products. Each gene product transcriptionally regulated a random set of other gene products in the network (with each gene having a 0.2 probability of regulating any other gene in the gene network). Gene self-regulation was allowed. Each regulation could be, with equal chance, either positive or negative (transcriptional activation and repression). Thus, every gene had, on average four connections, two positive and two negative connections,two efferent and two afferent. The value of this regulation between any pair of genes ($t_{ij}$), gene *i* and gene *j,* is a random, uniformly distributed, value between 0 and $t_{max}$. Taken together these $t_{ij}$ elements constitute the T matrix of transcriptional regulation, that is a representation of the gene network. See section 2.7, for a description of how is $t_{max}$ chosen.

For simplicity we only consider transcriptional regulation between genes, although EmbryoMaker can implement other kinds of interactions between gene products and other kinds of molecules.

We defined gene 1 as the "root" of the gene network and discarded all networks in which there was no activation-path connecting (i.e. a sequence of genes in which each gene positively regulates the next in the sequence), directly or indirectly, each gene to gene 1. This did not ensure that each gene in a network would be expressed, since each gene could also receive multiple negative regulation from other genes, it does give a chance to all genes to be expressed. Whether they do or not depends on the precise values in the T matrix and the dynamics of the particular developmental mechanism. Gene 1 does not diffuse between cells.

Each gene had a 50% chance of being either an extracellularly diffusible gene product (in here we call these growth factors) or an intracellular gene product. The latter had a 12.5% probability of being apically located, 12.5% of being basally located and 25% of being homogeneously located in the cell. We chose that gene 1 always directly activates a gene that can diffuse extracellularly.

2.1.2 *Diffusion coefficient:*

The diffusivity or diffusion coefficient of a molecule is a proportionality constant between the molar flux due to diffusion and the gradient in the concentration of the molecule. It depends on the molecules' weight, hydration and shape (larger for small roundish molecules). A different random value between 0 and $D_{max}$ was assigned to the extracellular diffusion coefficient of each gene product. In this case the random values were chosen from a logarithmic distribution, $D_i=10\ exp(log10(D_{min})+b*log10(D_{max}/D_{min})$, where b is chosen randomly from a uniform distribution with $0 < b <=1$ and $D_{min}=10^{-8}$. This ensures that diffusion coefficients of each order of magnitude are equally likely. The diffusion coefficient of the gene products that do not diffuse in the extracellular space are specified in the same way. See table 4 for the values of $D_{max}$ and $D_{min}$ and section 2.7 for a justification of these values.

2.1.3 *Degradation rate:*

All molecules in cellular or extracellular space end up being degraded, either specifically or non-specifically. Again the rate of this degradation depends on the molecules size, hydration and shape (as the steric accessibility to proteolytic enzymes). A different random value between $\mu_{min}$ and $\mu_{max}$ (with uniform distribution) was assigned to the degradation rate of each gene product. See table 4 for the values of $\mu_{min}$ and $\mu_{max}$ and 2.7 for a justification of these values.

2.1.4 Gene regulation of node mechanical properties and behaviors

When constructing a random developmental mechanism from a gene network, we randomly chose a set of gene products to affect randomly selected node properties or cell behaviors. Each gene product *i*, except gene 1, had a 50% chance of being chosen for that set. Each gene product can only affect one node property or cell behavior. Which one it would affect is chosen randomly among all node properties and cell behaviors. The value of such regulation by a given gene *k*, $e_{lk}$ for node property $l$ and $c_{mk}$ for cell behavior $m$, was chosen with the same logarithmic distribution than the diffusion coefficients. This ensured that values in each order of magnitude were equally likely. For the cell behaviors of cell division, apoptosis and epithelial-mesenchymal transitions we took an uniform distribution. The minimal and maximal values were $e_{l,min}$ and $e_{l,max}$ for node mechanical properties and $c_{m,min}$ and $c_{m,min}$ for cell behaviors. Each such limits took a specific value depending on the exact node mechanical property $l$ or cell behavior $m$ considered (see table 4). See section 2.7 for a justification of these values.

Each gene product could be chosen to be a membrane-bound adhesion molecule instead of affecting a node property or cell behavior. The probability of being an adhesion molecule was the same than that of affecting a node property (that is 1/(number of node mechanical properties + number of cell behaviors +1) ). In addition, we forced at least one random gene per network to be an adhesion molecule. The B matrix for each such developmental mechanisms was then a $n_a$ x $n_a$, where $n_a$ is the number of adhesion molecules in a given gene network. The values of the B matrix, this is the binding affinities between each adhesion molecule, were chosen randomly between $e_{ADH,min}$ and $e_{ADH,max}$. $e_{ADH,min}$ and $e_{ADH,max}$ are the same limits than for the node property $p^{ADH}$, that is the unspecific binding affinity between cells.

In addition, each random developmental mechanism included a constitutive activation of cell division and cell differentiation homogeneously throughout all cells. We chose to enforce that because cell divisions take place in basically every developing embryo and almost all developing organs. Cell differentiation causes cell behaviors to slow down over developmental time and is motivated by the widespread slowing down of growth during embryonic development. We chose to include this differentiation to represent the fact that, usually, cells become more determined to specific cell fates during development. The effect of this constitutive cell division and differentiation is mimicked in the model by adding a factor, in practice gene 11, that is expressed in each cell and whose concentration remains constant over time (its expression is not affected by any gene nor does it affect any gene). The strength of the constitute cell division and differentiation is determined by giving random values (between 0.5 and 0.75) to the corresponding elements of the E and C matrices, $e_{growth,11}$ and $c_{differentiation,\ 11.}$

##### 2.1.5. Initial conditions

We started all our simulations from the same simple initial conditions: a flat hexagonal sheet of epithelial of 126 cells and an underlying layer of 126 mesenchymal cells (see Figure S2). Each epithelial cell was represented by a cylindrical element, with an apical node and a basal node, and each mesenchymal cell by a single spherical node. At the initial condition gene 1 had a concentration of 1 in the most central epithelial cell and 0 elsewhere.

The initial values of mechanical properties were the same among cells in the initial conditions (see table 5). These values were chosen to be as biologically realistic as possible and to lead to no changes in the embryo themselves. This means that if these initial conditions are run under EmbryoMaker (without specifying a gene network) the embryo morphology does not change. See section 2.7 for a more detailed justification of the values chosen for these parameters.

##### 2.1.6 Simulation of each developmental mechanism:

Each random gene network and the node properties and cell behaviors it regulates we call a developmental mechanism. Each specific set of the T, E, C and B matrices constitute, thus, a different developmental mechanism. How the embryo changes over developmental time is something predicted from those by EmbryoMaker (e.g. we do not specify which should be the expression level of a gene at a given time and cell, this simply arises from the dynamics of the model).

EmbryoMaker can simulate models where each cell is made of several cylinders, for epithelial cells, and several spherical nodes, for mesenchymal cells. For simplicity, however, we chose to represent each epithelial cell by a single cylinder and each mesenchymal cell and the ECM by a single node, not just in the initial conditions but through all the simulation.

2.1.7. End of development simulation.

Each developmental mechanism simulation gave rise to a specific embryo morphology (as specific distribution of cells in 3D). During simulation time an embryo development was stopped if any of the criterion described in this section were met. Criteria *e* to *l* identify highly aberrant morphologies that were considered inviable and not further analyzed.

a) After 50 of developmental time units. This criterion was arbitrarily chosen to avoid prohibitively long times to simulate the ensemble and yet allow for morphologically complex embryos.

b) After 10 hours of computation time. This criterion was arbitrarily chosen to avoid prohibitively long times to simulate the ensemble and yet allow for morphologically complex embryos.

c) If more than 5000 nodes arise in an embryo. This criterion was arbitrarily chosen to avoid prohibitively long times to simulate the ensemble and yet allow for morphologically complex embryos.

d) If all cells become differentiated (differentiation implies that cells stop activating cell behaviors and as such stop moving).

e) If there was no cell movement over a longer period of developmental time. This criterion was chosen to avoid spending computation time in embryos that would lead to no change from the initial conditions. We empirically found out that no complex morphologies will arise unless cells start changing their positions from early on. Every hour of model time unit simulation, we calculated the displacement between the position of each node and its position one model time unit before (starting from model time unit 3). Simulations were stopped if the average displacement was smaller than 0.05 mdu or if the average displacement was smaller than 10% of the displacement recorded until that simulation time.

f) No morphogenesis until a certain time (5 developmental time unit), determined by a morphology's failure to grow perpendicularly to the cell sheet. Again, we empirically found that no complex morphologies will arise unless cells start changing there positions growing out from the x-y plane early on. Concretely, we stopped simulations if the z-coordinate of the highest epithelial centroid (centroid between apical and basal node) was less than 10% above the average of the z-axes of all epithelial cells.

In all the above criteria the resulting morphology may still be viable and included in our analysis. Some other criteria were used to stop wasting computational time in simulating embryos that were considered to be non viable.

g) When a morphology spread over more than 50 mdu. In other words, when the distance between minimal absolute x,y or z and maximal absolute x,y or z among cells was larger than 50 mdu. This occurred when many cells were dis-attached from the many body of the embryo.

h) If epithelial cells became too long. When either more than 20% of epithelial cells were longer than 3 times their equilibrium radius, $P^{EQD}$, or 10 times their length in the initial conditions.

i) Massive tissue disintegration. We considered this to be the case when more than 5% of the epithelial nodes were closer to a node of the opposite face (and from a different cell) than to the closest node of their own face (apical to basal and *vice versa*) . This occurred often as a result of insufficient cell adhesion or extreme epithelial torque forces.

j) Failed cell separation: When more than 1% of cells were closer than 0.05 mdu to the equilibrium radius ($p^{EQD}$) of another cell, we considered the tissue as unnatural and inviable. This was an artifact due to an unnatural parameter combination, usually very low repulsion and high adhesion.

k) If division rate exceeded a maximal division rate of one division for per unit model time. This occur in those rare cases in which many gene products are positively regulating cell division. We found that embryos dividing at extremely high rates will either explode or produce morphologies with an unrealistic overlap or packing of cells in space (as defined by the other criterion).

l) If it took more than 1200 seconds of computation time to simulate less than 0.1 model time units. This was usually a hallmark of highly aberrant morphologies and was, anyway, too inefficient computationally.

The first four criteria (a,b,c and d) clearly imposed a limit on the complexity of the embryo morphologies observed but imposing such limit is unavoidable given a finite amount of available computational time.

*2.1.8. Criteria for inviable embryos:*

Simulation stopped through criteria a, b, c, or d may still be considered inviable if any of the following criteria applied:

m) If more than 5% of the epithelial cells were flipped. An epithelial cell was considered to be flipped if its apical-basal polarity was inverted from that of its immediate neighboring epithelial cells.

n) Unrealistically overcrowded: if more than 5% of nodes had more than 24 neighboring nodes at close distance (less or equal than their equilibrium distance) or more than 15% of nodes with more than 12 neighboring nodes within this distance.

o) Disconnected epithelia: if the epithelium was broken into separate pieces. To check that, we measured the number of connected cell sets in each embryo. Two cells where considered as connected if there was some overlap between their adhesion radii ($p^{ADD}$). A connected cell set was then a set of cells in which one can go from any cell in the set to any other cell in the set through a sequence in which each cell is connected to the next cell in the sequence. We only counted sets that consisted of at least 3 cells. If an embryo had 3 of these sets or more it was considered to be broken.

p) Disconnected cells: if more than 1% of epithelial cells were disconnected from any other epithelial cell.

q) Broken epithelia: if more than 5% of epithelial cells had only one epithelial neighbor or more than 10% had less than 3 epithelial neighbors.

These criteria could also be used to stop simulations on the run but that would have been computationally inefficient.

#### 2.2. Signaling-only ensemble

Only morphologies with trivial morphologies were found in the broad ensemble. We ran 100,000 random developmental mechanisms and found very few morphologies differing from the initial conditions in a non-trivial way (such as simple size changes without changes in shape, noisy flat epithelial sheets and broken epithelia). We found that nearly all the gene networks were unable to change gene expression patterns over space. This means that most genes would be expressed only in the most central cell, in its immediate neighbors. or not at all. As a result most cells did not activate any cell behaviors and, thus, there were no changes in cell positions nor morphogenesis.

To circumvent this problem we made a simpler ensemble, the signaling-only ensemble, in order to identify gene networks capable of producing temporally stable changes in gene expression over space, as in a previous publication (Salazar-Ciudad *et al*, 2000). This ensemble considers only gene networks and cell communication through diffusible growth factors (we ran 20,000 networks). No cell behaviors or mechanical properties were considered and there was, thus, no cell movement. We then used the networks identified in such way to construct another ensemble, the signaling ensemble, in which genes from these networks regulate some randomly chosen node properties or cell behaviors at randomly chosen intensities (see section 2.3). The gene networks, degradation rates and diffusion coefficients were determined as in the broad ensemble.

*2.2.1 Construction of gene networks:* Just as in the broad ensemble, section 2.1.1

*2.2.2 Diffusion coefficient:* Just as in the broad ensemble, section 2.1.2

*2.2.3 Degradation rate:* Just as in the broad ensemble, section 2.1.3

*2.2.4 Gene regulation of node mechanical properties and behaviors:* There is no regulation of node mechanical properties or cell behaviors.

*2.2.5. Initial conditions*

To allow for faster simulations the initial conditions consisted of a one-dimensional row of 50 epithelial cells with gene 1 expressed in a steep gradient from the last cell in the row. All the networks that were identified to produce pattern formation were then simulated again on the initial conditions of the broad ensemble (a flat epithelium) to ascertain their pattern formation capacities.

2.2.6 End of development simulation. Each gene network was simulated for a maximum of 400 simulation time units, unless some of the other conditions above were met.

*2.2.7 Classifications of the resulting patterns*

A total of 20,000 different gene networks in the signaling-only ensemble were simulated for 10 developmental time units. Gene networks that did not lead to temporally stable and spatially heterogeneous gene expression patterns for at least one gene were discarded. The temporal stability was checked by eye. The remaining gene networks were classified in different categories based on the number (1-6, many), width (3 categories) of gene expression stripes and the existence of additional transient patterns (waves, oscillations), see Figure S20.

#### 2.3 Signaling ensemble

This ensemble is constructed by choosing gene networks in the previous ensemble and allowing some of their genes, to regulate node mechanical properties and cell behaviors. Only genes that show a non-trivial pattern, i.e. the non-homogeneous or not follow the gradient in the initial conditions (see 2.3.2.), can be chosen to regulate some cell behavior or node mechanical properties. This ensemble is, thus, just like building the broad ensemble from a specific set of gene networks that are known to produce temporally stable and spatially heterogeneous gene expression patterns.

Each gene network in the signaling ensemble was built by randomly choosing one gene network within one randomly chosen gene network category in the signaling-only ensemble. This ensured that gene networks from rare gene network categories (*e.g.* gene networks producing multiple stripe patterns of gene expression) were not picked less often than gene networks from more common gene network categories (*e.g.* gene networks producing only one-stripe patterns). Finally, randomly chosen genes where randomly assigned to regulate node mechanical properties and cell behaviors with random values, as in the broad ensemble.

*2.3.1 Gene regulation of node mechanical properties and behaviors:* As in the broad ensemble, section 2.1.4 but with some differences. Genes that will regulate cell behaviors and node mechanical properties are chosen from genes that show a stable a non-homogeneous pattern of expression. The selected genes will be able to regulate more than one cell behavior or node property. The number of cells behaviors and node mechanical properties regulated in a developmental mechanisms was determined by a binomial distribution B(10,0.5). Once we have the number of cell behaviors and node properties that will be regulated, we randomly chose which ones these will be. Finally, for each of these cell behaviors and node properties, a random gene with a non-homogeneous distribution will be chosen to regulate it.

*2.3.2 Initial conditions:* The initial conditions in this ensemble were the same than in the broad ensemble except for gene 1 that had a linear concentration gradient from one corner of the sheet to the center, both in the epithelium and in the mesenchyme, thus allowing for anterior-posterior polarity. Thus, the concentration of gene1 in cell *i* was

$$g(t=0)_{1i} = 1 - \phi\left(\frac{d_{ci}}{d_{cm}}\right) \quad (16)$$

Where $g(t=0)_{1i}$ is the concentration of gene 1 in cell *i* at the initial conditions and $d_{ci}$ is the distance between cell i and the cell that has the maximal concentration of gene 1 (the cell in the border) and $d_{cm}$ is the distance between this latter cell and the center (or centroid) of the initial conditions sheet.

*2.3.4 Simulation of each developmental mechanism:* As in the broad ensemble, section 2.1.6. In addition each developmental mechanism was simulated at least twice, each time with a different random seed. In other words, a different random sequence of random numbers was used to simulate noise in each simulation.

*2.3.5 End of development simulation:* As in the broad ensemble, section 2.1.7

##### 2.4 Autonomous ensemble

This ensemble consists of the same developmental mechanisms that the signaling ensemble but without cell signaling (i.e. all extracellularly diffusing gene products are replaced by transcriptional factors without modification of the gene network topology). In the initial condition one gene is expressed in all cells (gene 1) and the rest are not expressed.

##### 2.5 Autonomous biomechanics-only ensemble

In this ensemble genes do not transcriptionally regulate each other but regulate node mechanical properties and cell behaviors. Each developmental mechanism in this ensemble is based in a specific different developmental mechanism in the signaling ensemble. To construct a developmental mechanism we first define three genes, which are homogeneously distributed in all the nodes of the initial embryo. One of these genes is constantly being actively transported to the apical side and another one to the basal side, while the third gene remains homogeneous throughout the simulation. This way, some intracellular dynamics are conserved, but they arise simply from intracellular transport by active diffusion.

To assign cell behaviors and node mechanical properties to each of the three genes, we take into account which gene was regulating the cell behaviors in the corresponding developmental mechanism in the signaling ensemble. This way, cell behaviors and node mechanical properties that were regulated by a gene product that was actively being transported to the apical side or to the basal side of the epithelial cells, will be assigned to the genes in the autonomous ensemble that are also transported to the same side. All other genes are assigned to the gene that is neither apically or basally translocated. Each cell behavior and node mechanical property in each developmental mechanism is regulated with the same strength that in the corresponding developmental mechanism in the signaling ensemble.

This ensemble is just a way to explore the range of morphologies possible by just activating cell behaviors, or changing node mechanical properties, in an homogeneous way over space. It is important to note, however, that an homogeneous activation of cell behaviors, or spatially homogeneous change in node mechanical properties, does not imply a resulting homogeneous embryo morphology. In fact, it is often not the case, since cell behaviors and changes in node mechanical properties, can induce forces that spread over the embryo and can result, in their interaction with the embryo's margins and the responses to cells, in rather heterogeneous spatial patterns.

##### 2.6 Gradient autonomous ensemble

To explore to which extent the morphologies found in the signaling ensemble were due to the cell signaling or, merely, to the gradient in gene product 1 we build the gradient autonomous ensemble. This ensemble uses the same developmental mechanisms as in the signaling ensemble as well as the same initial conditions (gene product 1 expressed in the same gradient, see 2.3.2.) but no signaling as in the autonomous ensemble.

##### 2.7 Ranges of the model parameters:

###### *2.7.1 Maximal transcription regulation strength per gene, $t_{max}$*

We chose $t_{max}$ to be such that the maximal rate of change of transcription rate in respect to $t_{ij}$ should be less than 0.1% in the simple case of a gene activating its own expression. The rate of transcription per unit time of gene $i$, $Q_i$, in the case that it regulates itself and is not regulated by any other gene is:

$$Q_i = \frac{t_{ii}g_i}{(t_{ii}g_i + 1)} \qquad (17)$$

The derivative of $Q_i$ in respect to $t_{ii}$ is then:

$$\frac{\partial Q_i}{\partial t_{ii}} = \frac{g_i}{(t_{ii}g_i + 1)^2} \qquad (18)$$

If run long enough the concentration of $g_i$ will reach an equilibrium. At equilibrium the time derivative of $g_i$ is zero:

$$\frac{\partial g_i}{\partial t} = \frac{t_{ii} g_i}{(t_{ii} g_i + 1)} - \mu_i g_i \qquad (19)$$

$$\hat{g}_i = \frac{1}{\mu_i} - \frac{1}{t_{ii}} \quad (20)$$

If the initial condition of $g_i$ is larger than this equilibrium concentration then $g_i$ will decrease over time until it becomes equal to this equilibrium concentration. If the initial $g_i$ is smaller than that then the equilibrium concentration would also be the maximal concentration, $g_{max}$.
Inserting (20) into (19) and deriving again with respect to $t_{ii}$ we obtain:

$$\frac{\partial^2 Q_i}{\partial t_{ii}^2} = \mu_i^2 \frac{\mu_i t_{ii} + \mu_i - t_{ii}}{\mu_i (t_{ii} - 1) + t_{ii})^3} \qquad (21)$$

We chose $\mu_i$ to be such that, for the case of a gene regulating itself, the equilibrium concentration $\hat{g}_i$ does not decrease by more than 0.1% when we increase $\mu_i$. This is:

$$\frac{\partial \hat{g}_i}{\partial \mu_i} = -\mu_i^{-2} = 0.001 \quad \text{and then} \quad \mu_i = 31.62 \quad (22)$$

Replacing this value in equation 21 and demanding that $\frac{\delta^2 Q_i}{\delta t_{ii}^2} = 0.001$ we obtain that the maximal $t_{ii}$, now $t_{max}$, is 31.62.

*2.7.2 Diffusion coefficient:*

To define a meaningful range of extracellular diffusion coefficient for gene products, we considered that in the ensemble the average cell size was 0.25 *mdu* (model distance units) and in a real epithelial tissue 30 μm. Then each model distance unit represents 120 μm. The extracellular diffusion coefficient of ovalbumin is 0.675 $cm^2/s$ (Culbertson *et al*, 2002), into model units that is a diffusion coefficient of $D$=0.021 $mdu^2/s$. Since diffusion rates are available only for very few proteins, we considered ovalbumin as a well-studied and relatively fast diffusing protein and defined the minimal diffusion rate to be two orders of magnitude below and the maximal diffusion rate to be two orders of magnitude above this value.

*2.7.3 Degradation rate:*

From equation (8) it can be seen that the maximal rate of production of any gene product is 1 (since the sum of transcriptional regulations from other genes is both in the numerator and denominator). From equation (8) it can also be deduced that any gene product concentration will reach, as time progresses, an equilibrium value of $g_{max}=1/\mu$ . From zero concentration initial conditions (or from any initial condition with a gene concentration less than $1/\mu$) this equilibrium concentration is also the maximal concentration any gene product can reach. From this condition it follows that this maximal gene product concentration decreases with $\mu$ but does it very slowly when $\mu$ is large. In fact, it goes as $d\hat{g}/d\mu = -\mu^{-2}$. We chose $\mu$ to be such that this rate of change would be smaller than 0.1%. This is $\mu_{max}=32$. We chose $\mu_{min}$ such that the maximal concentration of gene products within a cell will never be larger than one, following $g_{max}=1/\mu$, this is $\mu_{min}=1$. From that it follows that the largest concentration a gene can reach in the ensemble is $g_{max}=1/\mu_{min}$, that is 1.

*2.7.4* Regulation of node properties, $e_{k,min}$ and $e_{k,max}$:

As described in section 1.5 node properties can change over time due to changes in gene expression, equation (12). How strongly a gene *k* can regulate a node property is determined by each $e_{lk}$ element in the E matrix, where *l* is a node property. In this section we describe how we chose the maximal and minimal values, $e_{k,min}$ and $e_{k,max}$, for these elements.
In the developmental mechanisms in our ensembles most node properties are regulated by only one gene product. In this simple case the maximal possible value of a node property *l* during a simulation would be $p^l_{max} = p^l(0) + g_{max}\, e_{l,max}$. Since $g_{max} = 1$ then $p^l_{max} = p^l(0) + e_{l,max}$ and $e_{l,max}$ simply represents how much a node property increases from its value in the original conditions.

Most gene networks chosen from the signaling-only ensemble to construct the signaling ensemble, expressed genes at very different concentrations. Since we calculated the range of strengths at which genes would regulate cell behaviors (C matrix) and node biomechanical properties (E matrix) based on the maximum possible gene product concentration ($g_{max}$=1), we decided to correct for those differences. Otherwise a weakly expressed gene would never be able to regulate a cell behavior at high rates. Thus, we divided, for each gene *k* in each developmental mechanism, all the $e_{lk}$ and $c_{nk}$ values by the average concentration of *k* over time in the cells where it was expressed in the signaling-only ensemble. Then for each network in the signaling ensemble we calculated the average concentration of gene product *k* in 10 different simulations (each using a different random seed) in which node properties and cell behaviors where regulated by these normalized $e_{lk}$ and $c_{nk}$ values for each gene. This was necessary, since in the signaling signaling ensemble the actual gene product concentration also depended of cell size, geometry and cell movements and duration of the simulation. The original values of $e_{lk}$ and $c_{nk}$ where then divided by this latter average expression value for each gene *k*.

2.7.4.1 Components of $p^{EQD}$: $p^{COD}$, $p^{GRD}$, $p^{PLD}$ and $p^{VOD}$:

In animals, epithelial cell diameters range between 1 and 100 µm (Alberts *et al*, 2002) and average around 30 µm. The model equivalent of cell diameter would be 2 times the equilibrium radius, $p^{EQD}$. This latter radius is 0.25 mdu in the initial conditions and, thus, each mdu corresponds to 60 µm. The maximal epithelial diameter of 100 µm is then 1.67 mdu. We demanded that $p^l_{max}$ would be smaller than half the maximal cell length observed in animals, 0.83 mdu. We chosen $e_{l,max}$ to be equal to that $e_{l,min}$ to be two orders of magnitude below.

2.7.4.3. *$p^{ADD}$, adhesion radius:*

$p^{ADD}$ is the equilibrium radius plus the distance at which a cell can extent projections (filopodia) to make cell contact. Long filopodia are reported to be 0.125md (Nilufar *et al*, 2013), so we added this value to the average equilibrium range. Since measurements on filopodial growth are scarce, we cannot exclude substantially larger filopodial lengths. Thus, for $e_{ADD,max}$, we add an order of magnitude and reduce two for $e_{ADD,min}$. Thus, $e_{ADD,min} = 0.0375$ and $e_{ADD,max} = 3.75$.

*2.7.4.4. $p^{YOU}$: intracellular elasticity:*

The elasticity parameter $p^{YOU}$ determines the force that binds together two nodes that are at a distance smaller than the sum of their adhesion radii. From equation (2) the elasticity force acting between two nodes *i* and *j* is:

$$f_{Aii} = k_{ij}^{ADH}(d_{ij} - p_i^{EQD} + p_j^{EQD}) \quad (23)$$

In the absence of other forces and due to the over-damped nature of our equations the rate of displacement of node i over time would be equal to that force.
Solving the temporal differential equation, we obtain a formula of displacement as a function of time:

$$r_i(t) = r_{io}^{-2p^{YOU}t} \qquad (24)$$

From biophysical experiments (Farhadifar *et al*, 2007) the relaxation half-time of an elastic tissue can be estimated as 6.5s. By rotating the *X* coordinate axis to be in the line joining nodes *i* and *j, r* becomes *x*. Then x(t+6.5s) = 0.5 x(t) and

$$p^{YOU} = \frac{ln(0.5)}{-2 \times 6.5} = 0.0533 \qquad (25)$$

Since biomechanical experiments are scarce, we considered the calculated value of $p^{YOU}$, based on experiments, as a biological average and set minimal and maximal values two orders of magnitude above and below this value. Thus $e_{YOU,min} = 0.00053$ and $e_{YOU,max} = 5.3$.

2.7.4.5. $p^{REC}$, cell compressibility, $p^{ADH}$, intercellular adhesion, $p^{HOO}$, apico-basal elasticity and the epithelial torsion properties $p^{ERP}$ and $p^{EST}$. All these node properties ultimately define a rate of tissue relaxation, we use therefore the same $e_{l,max}$ and $e_{l,min}$ than with $p^{YOU}$.

2.7.4.6. $p^{EQS}$, Apico-basal equilibrium distance. $e_{EQS,max}$ and $e_{EQS,min}$ where chosen to be equal to $e_{COD,max}$ and $e_{COD,min}$ values since the later are related to how large cells can be while the formed are related to how long epithelial cells can be. These latter values should be relatively similar, at least in the same order of magnitude, and we thus make them the ranges that depend on them, $e_{EQS,max}$ and $e_{EQS,min}$ , equal.

2.7.4.7. $p^{MOV}$, Filopodial instability: The likelihood that an energetically unfavorable node movement is nevertheless accepted. We keep $e_{MOV,min}$ and $e_{MOVmax}$ between 0.0001 and 10.

2.7.4.8. $p^{DMO}$, Filopodia extensibility: We use the average cell diameter including filopodia, which is twice $e_{ADD,max}$. Thus, $e_{DMO,min}$=0.075 and $e_{DMO,max}$=7.5

2.7.4.9. $p^{DIF}$, cell differentiation. We defined minimum and maximum so that a given cell would fully differentiate between 10 and 100 model time units. This is the case if $e_{DIF,min}$=0.075 and $e_{DIF,max}$=0.1

*2.7.5 Regulation of cell behaviors, $c_{k,min}$ and $c_{k,max}$:*

The regulation of many cell behaviors takes the general form:

$$\frac{\partial P_i^B}{\partial t} = \frac{1}{n_i} \sum_{i=1}^{n_i} \sum_{m=1}^{n_g} c_{m,B} g_{im} \qquad (26)$$

Where *B* is the cell behavior variable in cell *i* . $n_i$ is the number of nodes in cell *i* (1 per mesenchymal cells and two for epithelial cells) and $n_g$ the number of genes in a network. $c_{m,B}$ is the value of the regulation of cell cycle by gene product *m* and $g_{im}$ is the concentration of gene product *m* in cell *i*. Since most cell behaviors are regulated by just one gene product and $g_{max}$=1 it follows that:

$$\frac{\partial P_i^B}{\partial t} = c_{m,B} \qquad (27)$$

The definite integral of this expression over one model time unit (between 0 and 1) is just $c_{m,B}$. We then chosen these values based on the maximal number of cell behaviors events (e.g. cell division) per cell per model unit time .

*2.7.5.1. Cell division:*

Cell's *i* cycle phase, cell property $P_i^{PHA}$, is a continuous variable between 0 and 1. When this variable reaches 1 the cell divides and the variable takes value 0, and then can grow again. $c_{k,max}$ was chosen so that at most cells will divide once per developmental time unit (that is once per hour). Faster cell divisions leads to artefactual tissue behavior, as cells would undergo multiple divisions before having enough time to separate from their sister cells. Then to get at maximum of 1 division per model unit time $c_{m,PHA} = c_{PHA,max}=1$. $c_{PHA,min}$ is just 2% of this value.

*2.7.5.2. Apoptosis:*

Cell death has been reported to require at least approximately 20 times as long (Kerr *et al*, 1972) as the fastest cell divisions rounds (O'Farrell *et al*, 2004). Based on values chosen for cell division we chose $c_{APO,max}=0.005$ and $c_{APO,min}=0$.

*2.7.5.3 Epithelium-to-mesenchyme transition (EMT):*

EMT, for instance evident in the appearance of primary mesenchymal cells during the sea urchin cleavage, has been described to take approximately twice as long as the fastest cell divisions, which we use as an upper limit (Wu *et al*, 2007). Thus, $c_{EMT,max}=0.05$ and $c_{EMT,min}=0$.

*2.7.5.4. ECM secretion:*

Since we do not define the nature of the ECM units secreted here, we limit secretion so that, if 100 cells in a morphology keep secreting at maximal secretion rate, the morphology does not accumulate more than about 5000 nodes (including cells and ECM units) before at least reaching 1 developmental time unit. The lower limit is the same as for division. $c_{ECM,max}=0.05$ and $c_{ECM,min}=0$.

ECM nodes have their own node properties, similar to mesenchymal nodes. The properties of the ECM nodes are established by the gene regulating the production of ECM. The limits of the node properties are the same as for the other nodes types. The exception is , which for ECM nodes it has a maximum value of 1.0.

#### 2.8. Node mechanical property values in the initial conditions:

Table 5 shows these initial values in the initial conditions. The radii at which two nodes will start to experience a repulsion force, $p^{EQD}$, or an adhesion force, $p^{ADD}$, were chosen to be 0.25 mdu and 0.35 mdu respectively. These numerical values define the spatial scale of the model. The proportion between these two values, however, defines how much a cell can be compressed before experiencing a restoring force. In a way $p^{ADD}$ can be understood as the maximal distance (from the cell center) at which non-migrating cells can extent adhesive cytoplasmatic projections. Many of the values of node mechanical properties are chosen based on these two fundamental values.

The values of many node mechanical properties were chosen so that, if unchanged, the embryo would grow without breaking. This is the case for: $p^{REP}$ and $p^{REC}$, the repulsion force constants (these take the same value in the case in which cells are made of single element, as in the ensemble), $p^{YOU}$ and $p^{ADH}$, the adhesion force constant (that are the same in the case in which cells are made of single element), the equilibrium distance between the apical and basal node of a cylinder, $p^{EQS}$, the elasticity constant of the apical-basal spring-link for each cylinder, the resistance to bending of the epithelium, $p^{ERP}$, and the stability and extensibility of cell cytoplasmatic projections, $p^{MOV}$ and $p^{DMO}$. The rest of node mechanical properties were set to zero.

**3.** *Orientation Morphological Distance (OMD)*

*3.1 Orientation Morphological Distance (OMD).* Given morphologies A and B, we first calculate the orientation of each cell in A and B (just as in the OPC measurement). As a result, each cell has an integer value between 0 and 8, depending on their orientation. Second, we stretch the epithelium of A and B, making it flat (2D), (see SI 3.2 for more details on how we do this). Third, we center morphology A and B at their respective cell one and then superimpose A and B. Forth. Each cell in morphology A will be compared to the closest cell in morphology B and each cell in B to the closest cell in A, if the compared cells have different orientation, we add one to the OMD distance. Finally, we divide OMD by the total number of cells in morphology A and B. The OMD distance will range from 0 to 1, where 1 indicates that the morphologies are completely different.

3.2. Flattening of the epithelium

To flatten the epithelium, epithelial cells were ordered according to the length of the minimal neighbor path to node 1 ($l_{mnp}$). Nodes in the epithelium rarely interchange neighbors. so this node is always located around the center of the embryo. A neighbor path is a list of cells in which each cell in the list is in physical contact with the next cell in the list and in which cell 1 is the last element of the list. It is essentially a path of cells from which one could go from a cell to cell 1, by jumping between cells that are in physical contact. The minimal neighbor path is the shortest (i.e. measured as order, or number of neighbors. in the list) possible list between a cell and cell 1. For each cell we also calculated the accumulated angle, in the plane of the epithelium, they have to rotate along the minimal neighbor path to reach cell 1. Based on the minimal neighbor path and that angle, each cell gets a position in the XY plane that closely resembles the one it would have had if the epithelium would be stretch until it completely flattens. For each embryo we calculated the cell with the largest minimal neighbor path (roughly the cell that is more far away from cell 1 in the plane of the epithelium). Only cells with a minimal neighbor path of a length shorter than 2/3 of minimal neighbor path of this cell were considered to calculate the OMD (the shortest of this largest path among the two embryos compared was taken). This was done to avoid effects due to the irregular margins of the epithelium. The position of each cell *i* in the XY plane was thus:

$$x_i = l_{mnp}\ cos\left(\alpha_a\right)$$
$$y_i = l_{mnp}\ sin\left(\alpha_a\right)$$

Where $\alpha_a$ is the accumulated angle.

**4. Other analysis.**

*4.1 Disparity*

To calculate the disparity of one morphology, we calculated the EMD distance to all other morphologies in the same ensemble, the disparity value is the average of these distances. Due to the high number of morphologies in each ensemble, we calculated the disparity only for a subset of each ensemble. Each subset is made of 500 randomly chosen morphologies. Only morphologies whose complexity by angle variation complexity was higher than 0.3 were included in the plots.

*4.2 Developmental stability*

The stability of a developmental system is calculated as the EMD distance between two twins. The morphologies are centered in order to avoid introducing an additional source of noise. Due that the initial conditions of all embryos is the same is not necessary to rotate the embryos to find the least EMD distance between two morphologies.

##### 4.3 **Number of cells contracting in the same way**

For all morphologies in the signaling ensemble we calculated the number of cells that have a similar radius and are in contact to each other as forming a single patch (six groups are made, from 0.1 to 0.5 in 0.1 intervals). This was done for epithelial cells, taking into account apical and basal nodes separately. Then we calculate the size of the largest of such patches in each morphology. Only morphologies with a total size of at least 1000 epithelial nodes that have at least two patches and whose complexity by angle variation complexity was higher than 0.3 were included

#### 5. References:


1. Marin-Riera M, Brun-Usan M, Zimm R, Välikangas T, Salazar-Ciudad I (2015) Computational modeling of development by epithelia, mesenchyme and their interactions: a unified model. *Bioinformatics* 32(2):219-25.
2. Brun-Usan M, Marín-Riera M, Grande C, Truchado-Garcia M, Salazar-Ciudad I (2017) A set of simple cell processes is sufficient to model spiral cleavage. *Development* 144(1)
3. Salazar-Ciudad I (2003) Mechanisms of pattern formation in development and evolution. *Development* 130(10):2027–2037.
4. Graner F, Glazier JA (1992) Simulation of biological cell sorting using a two-dimensional extended Potts model. *Phys Rev Lett* 69(13):2013–2016.
5. Salazar-Ciudad I, Garcia-Fernández J, Solé RV (2000) Gene networks capable of pattern formation: from induction to reaction-diffusion. *J Theor Biol* 205(4):587–603.
6. Salazar-Ciudad I, Newman SA, Solé RV (2001) Phenotypic and dynamical transitions in model genetic networks. I. Emergence of patterns and genotype-phenotype relationships. *Evol Dev* 3(2):84–94.
7. Reinitz J, Sharp DH (1995) Mechanism of eve stripe formation. *Mech Dev* 49(1–2):133–58.
8. Culbertson CT, Jacobson SC, Michael Ramsey J (2002) Diffusion coefficient measurements in microfluidic devices. *Talanta* 56(2):365–373.
9. Alberts B, et al. (2002) *Molecular biology of the cell* (Garland Science).
10. Nilufar S, Morrow AA, Lee JM, Perkins TJ (2013) FiloDetect: automatic detection of filopodia from fluorescence microscopy images. *BMC Syst Biol* 7:66.
11. Farhadifar R, Röper J-C, Aigouy B, Eaton S, Jülicher F (2007) The Influence of Cell Mechanics, Cell-Cell Interactions, and Proliferation on Epithelial Packing. *Curr Biol* 17(24):2095–2104.

12. Kerr JF, Wyllie AH, Currie AR (1972) Apoptosis: a basic biological phenomenon with wide-ranging implications in tissue kinetics. *Br J Cancer* 26(4):239–57.

13. O'Farrell PH, Stumpff J, Su TT (2004) Embryonic cleavage cycles: how is a mouse like a fly? *Curr Biol* 14(1):R35-45.

14. Wu S-Y, Ferkowicz M, McClay DR (2007) Ingression of primary mesenchyme cells of the sea urchin embryo: A precisely timed epithelial mesenchymal transition. *Birth Defects Res Part C Embryo Today Rev* 81(4):241–252.

#### Supplementary figures

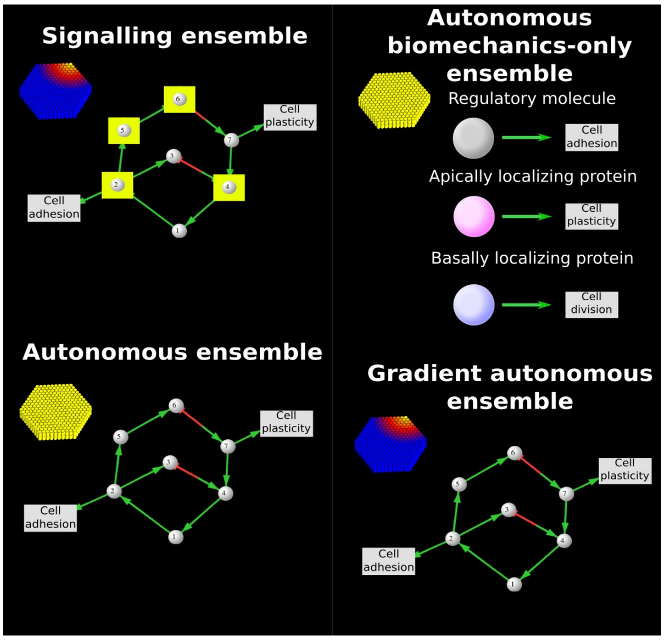


**Figure S1. Example developmental mechanisms and pattern transformations.** The figures shows two identical initial developmental patterns, on the left, and two different resulting developmental patterns, to the right. Each developmental pattern is a specific distribution of cell types in space. The networks in the center are two developmental mechanisms that are responsible from the transformation of the initial developmental patterns in the left to the resulting developmental patterns in the right (as simulated in our EmbryoMaker model). In the initial developmental patterns, cylinders represent epithelial cells and their color represents the level of expression of a gene (yellow maximal value, blue minimal value). In the resulting developmental patterns colors show the value of the position of each cell in the z-axis. In the developmental mechanisms circles represent gene products and arrows regulatory interactions (positive in green and negative in red) either on other gene products or on cell behaviours or node mechanical properties (these latter are shown in grey boxes). The circles surrounded by yellow boxes indicate growth factors (gene products that diffuse in the extracellular space and can then affect gene expression in other cells other then where they are expressed).

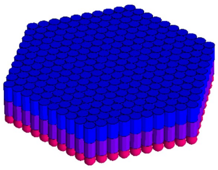

**Figure S2. Initial developmental pattern.** The figure shows the basic initial conditions used in all simulations. Initial gene expression patterns are not shown since they differs between the ensembles. Cylinders represent epithelial cells and spheres mesenchymal cells. Each cylinder is made of two elements, a basal one (in blue) and an apical one (in violet). In total, there are 271 epithelial cells and 271 are mesenchymal cells.

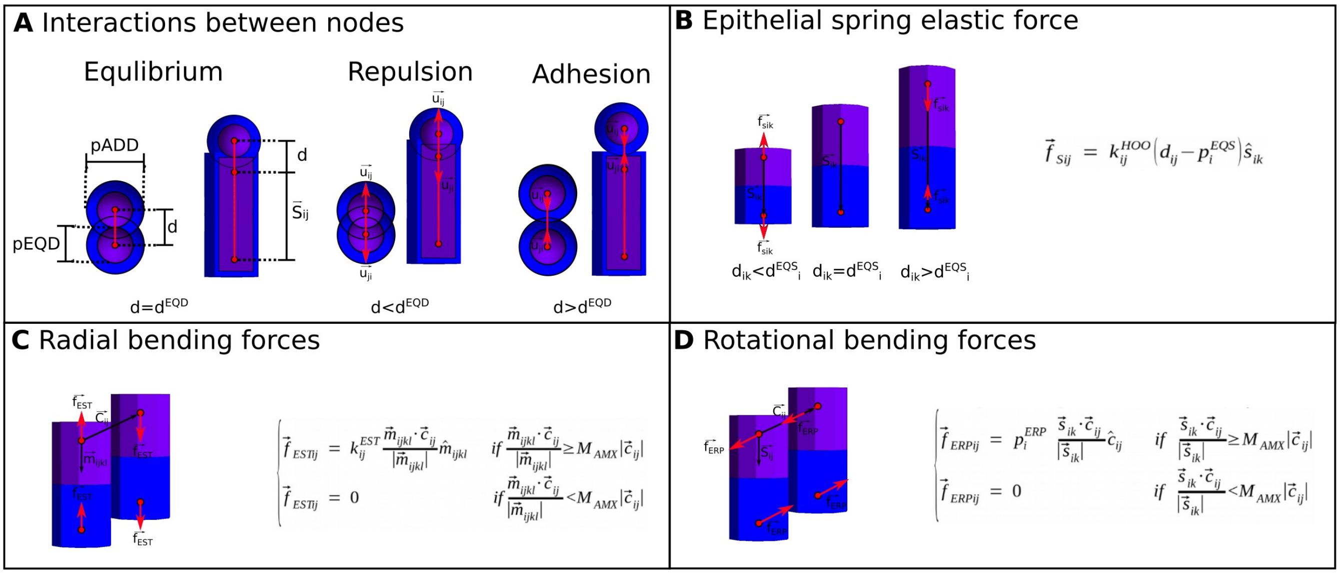


**Figure S3. Basic mechanical interactions in EmbryoMaker. A.** Mechanical interactions between spherical nodes and between spherical and cylindric nodes are determined by the distance between their centers and their distance of equilibrium. When two spherical nodes (either mesenchymal or ECM) are at a distance (d) smaller than $d^{ADD}$ they experience an attractive force, when they are closer than $d^{EQD}$ they experience a repulsive force. $d^{EQD}$ is simply the sum of the sizes or radius of repulsion, $p^{EQD}$, of the nodes interacting while $d^{ADD}$ is the sum of the radius of adhesion of the nodes interacting, $p^{ADD}$. Notice that these radii are variables of the model and then can changes over time as a result of gene expression or external pressure changes. The direction of the force (red arrows) goes from the center of one node to the center of the other. When a spheric node interacts with a cylinder's apical or basal face, the direction of the force is always parallel to the apical-basal axis of the cylinder (and perpendicular to it when the interaction is lateral). **B, C and D** are mechanical forces specific for epithelia. **B.** The two nodes composing a cylinder are connected by an unbreakable spring (black line). Elastic forces will always follow the direction of that spring. The spring has an equilibrium distance ($d^{EQS}$), if the distance between the centers of the nodes (d) in a epithelial cylinder are closer than $d^{EQS}$ (left figure in B), elastic forces will repulse the nodes (red arrows). If the distance between the centers of the nodes is greater than $d^{EQS}$, elastic forces will attract them (this distance is again the sum of the mechanical property, $p^{EQS}$, of the two nodes that is itself a variable of the model. **C.** Epithelial bending forces tend to put two cylinders in a position in which the angle between the vector connecting the two apical (or basal) nodes and the apical-basal axis is $\pi/2$. Epithelial bending forces applies on a direction normal to the apical/basal surface. **D.** The bending rotational force applies in the direction connecting the two epithelial nodes from the same side.

| Embryo | AV | OPC | Embryo | AV | OPC |
|---|---|---|---|---|---|
| 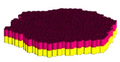 | <0.001 | 1 | 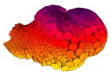 | 0.73 | 19 |
| 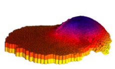 | 0.13 | 6 | 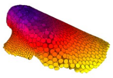 | 1.25 | 19 |
| 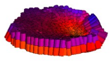 | 0.23 | 11 | 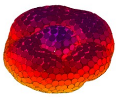 | 1.41 | 30 |
| 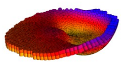 | 0.33 | 14 | 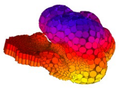 | 1.98 | 68 |
| 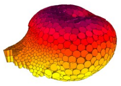 | 0.53 | 23 | 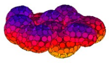 | 2.67 | 108 |

**Figure S4. Examples of embryos from the ensembles sorted by their complexity.** The color of each embryo shows the relative position of the nodes in the Z-axis. For each embryo the complexity is show as angle variation (AV) and as orientation patch count (OPC).

| Embryo 1 | Embryo 2 | Developmental instability (EMD) | Developmental instability (OMD) | Embryo 1 | Embryo 2 | Developmental instability (EMD) | Developmental instability (OMD) |
|---|---|---|---|---|---|---|---|
| 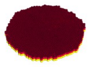 | 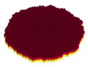 | 0.065 | 0.006 | 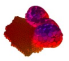 | 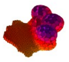 | 0.235 | 0.477 |
| 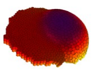 | 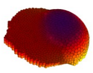 | 0.085 | 0.021 | 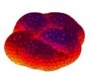 | 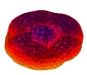 | 0.255 | 0.583 |
| 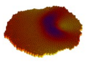 | 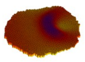 | 0.115 | 0.061 | 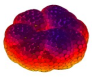 | 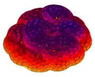 | 0.260 | 0.600 |

**Figure S5. Examples of embryos from the ensembles sorted by their developmental instability.** The figure shows in each row a pair of simulations of a developmental mechanisms. Due to noise these two resulting morphologies can differ. The third column shows a measure of the morphological distance between them (measured as EMD) and the fourth column shows the morphological distance measured as OMD . The color of each embryo's morphology shows the relative position of the nodes in the Z-axis.

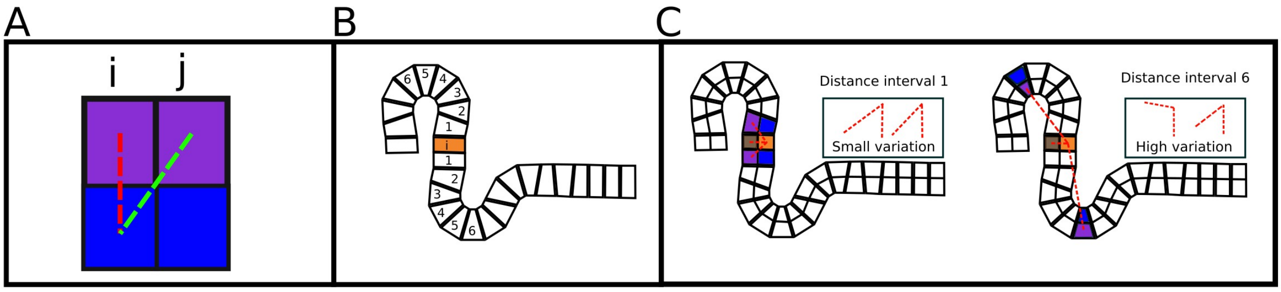


**Figure S6. Angle-distance variance. A.** The angle between two cells is calculated using two vectors. One apical-basal vector (in red) and one apical-apical vector (in green). **B.** Cells are classified in categories depending on the distance to cell *i*. **C.** The variation of angle variation inside each distance interval is calculated.

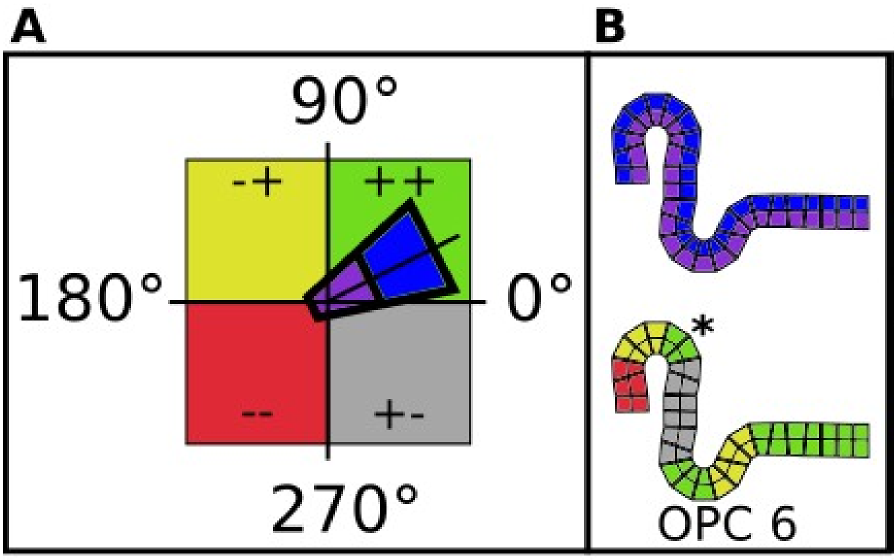


C

| Octal | X | Y | Z |
|---|---|---|---|
| 1 | - | - | - |
| 2 | + | - | - |
| 3 | - | + | - |
| 4 | + | + | - |
| 5 | - | - | + |
| 6 | + | - | + |
| 7 | - | + | + |
| 8 | + | + | + |

**Figure S7. Orientation patch count.** With OPC groups of contiguous cells with the same orientation are grouped together into a patch. In A we can see how using the apical and basal nodes of the epithelial cells we can determine the cells orientation. The orientation of the cells is categorized as in C. In B we can see how patches are formed in an epithelium, notice that we have 7 patches in the bottom figure in B, but only 6 are counted towards the OPC value since the patch pointed out by the asterisk is too small to be taken into account.

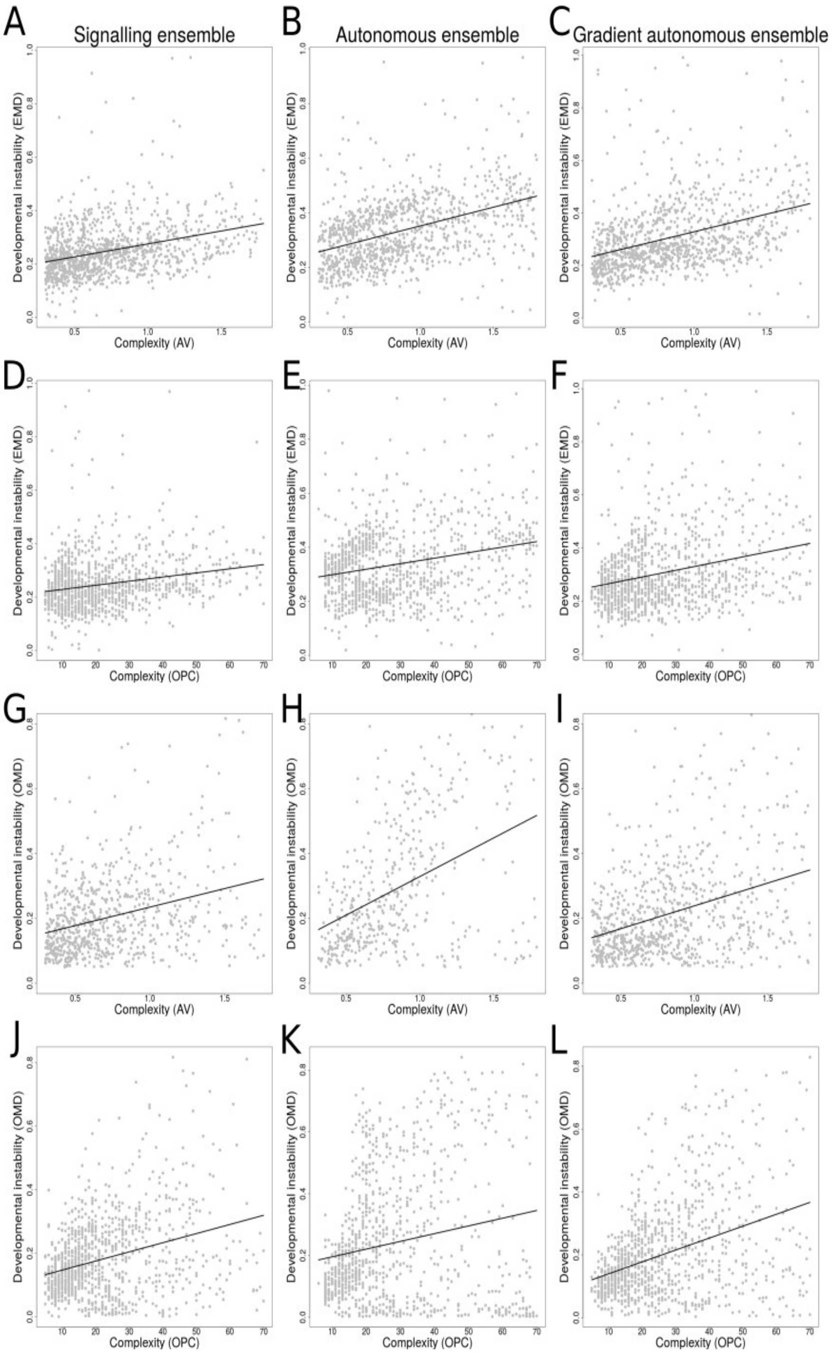


**Figure S8. For morphologies of a given complexity developmental instability is lower in the narrow and gradient ensembles.** Plots show, for each morphology in the three ensembles, complexity on the x-axis and developmental instability on the y-axis. Notice that across morphologies of different complexities those from the signaling ensemble tend to be more developmentally stable than morphologies in the autonomous gradient ensemble and the autonomous ensemble. The plots that use AV complexity, only show the results for AV higher than 0.3, the plots that use OPC, only show the results for OPC higher than 5. The top six plots use EMD as the measurement for developmental instability, while the six lower ones, use OMD as the developmental instability measurement. The black lines shows the lineal regression. **A**. Spearman correlation: $r_s$=0.416, pval<0.001, n=804 . **B.** Spearman correlation: $r_s$=0.418, pval<0.001, n=508. **C.** Spearman correlation: $r_s$=0.425, pval<0.001, n=731. **D.** Spearman correlation: $r_s$=0.237, pval<0.001, n=810. **E.** Spearman correlation: $r_s$=0.022, pval=0.625, n=478. **F.** Spearman correlation: $r_s$=0.218, pval<0.001, n=821. **G.** Spearman correlation: $r_s$=0.297, pval<0.001, n=729. **H.** Spearman correlation: $r_s$=0.430, pval<0.001, n=428. **I.** Spearman correlation: $r_s$=0.337, pval<0.001, n=776. **J.** Spearman correlation: $r_s$=0.404, pval<0.001, n=731. **K.** Spearman correlation: $r_s$=0.469, pval<0.001, n=419. **L.** Spearman correlation: $r_s$=0.497, pval<0.001, n=768. The pvalues for the Spearman correlation was calculated with a permutation test (9999 repetitions). All ensembles have significantly different distributions (pval<0.001, tested with Kruskal-Wallis). **A vs B**: pval<0.001, H=163.415, n=1312. **A vs C**: pval<0.001, H=83.114, n=1535. **B vs C**: pval<0.001, H=20.233. **D vs E**: pval<0.001, H=156.437, n=1288. **D vs F**: pval<0.001, H=77.376, n=1631. **E vs F**: pval<0.001, H=21.360, n=1299. **G vs H**: pval<0.001, H=57.612, n=1157. **G vs I**: pval<0.001, H=12.194, n=1505. **H vs I**: pval<0.001, H=23.036, n=1204. **J vs K**: pval<0.001, H=66.418, n=1150. **J vs L**: pval<0.001, H=12.598, n=1499. **K vs L**: pval<0.001, H=25.225, n=1187.

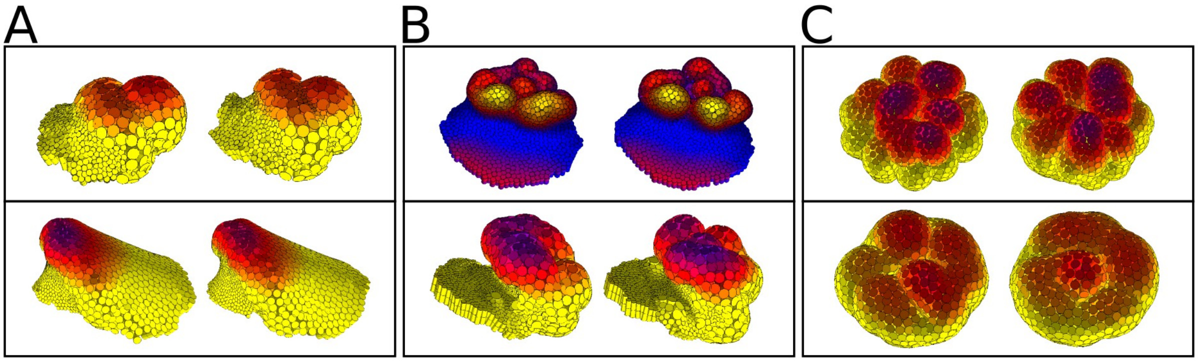


**Figure S9. Examples of complex morphologies with different developmental instability**. **A.** Complex and stable embryos. In these embryos the invaginations appear in the same position in different runs. Top row. Developmental instability: EMD=0.1, OMD=0.14; Complexity (AV): 0.74; Complexity(OPC): 18. Bottom row. Developmental instability: EMD=0.15, OMD=0.13; Complexity (AV): 0.79; Complexity(OPC): 11. **B.** In these embryos only half of the morphology undergoes significant morphogenesis, they are complex, but not very stable. Top row. Developmental instability: EMD=0.11, OMD=0.18; Complexity (AV): 1.77; Complexity(OPC): 62. Bottom row. Developmental instability: EMD=0.18, OMD=0.42; Complexity (AV): 1.89; Complexity(OPC): 72. **C.** Very complex and very developmentally unstable morphologies. In these morphologies invaginations appear in different locations every time the same developmental mechanism is run. Top row. Developmental instability: EMD=0.38 , OMD=0.79; Complexity (AV): 2.54; Complexity(OPC): 162. Bottom row. Developmental instability: EMD=0.25, OMD=0.583; Complexity (AV): 1.35; Complexity(OPC): 42

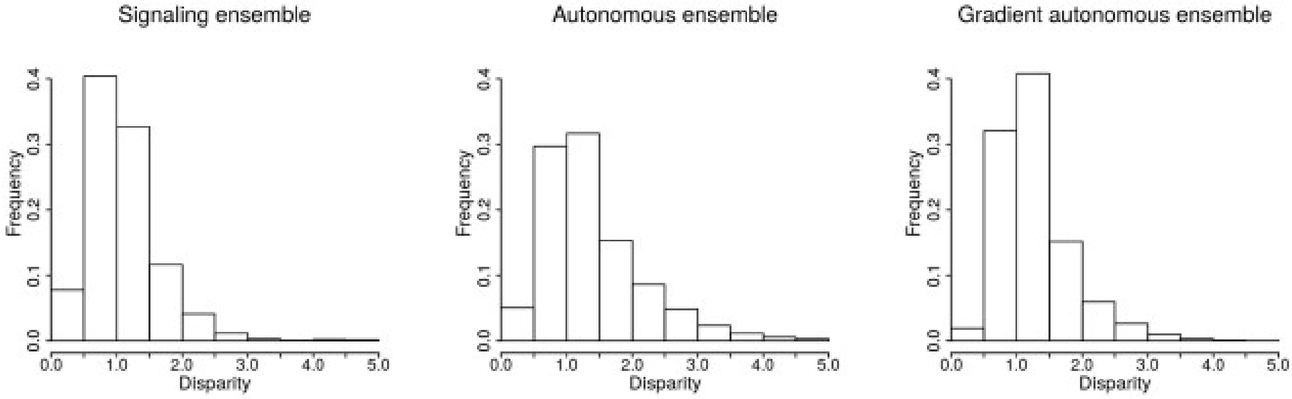


**Figure S10. Cell-cell signaling does not increase overall morphological disparity.** Frequency distributions for morphologies of three ensembles, sorted by morphological disparity. In each plot, the x-axis shows the average disparity per morphology, while bar heights correspond to relative frequencies (100% = 1). To calculate the disparity of one morphology, we calculated the EMD distance to all other morphologies in the same ensemble, the disparity value is the average of these distances. Due to the high number of morphologies in each ensemble, we calculated the disparity only for a subset of each ensemble. Each subset is made of 500 randomly chosen morphologies. Only morphologies whose complexity by angle variation complexity was higher than 0.3 were included in the plots. We did a Wilcoxon-Mann-Whitney test to study the differences between the ensembles. Signaling ensemble and autonomous ensemble: pval<0.001, Z=102.764, n=124750. Signaling ensemble and gradient autonomous ensemble: pval<0.001, size effect

Z=89.482, n=124750 . Autonomous ensemble and gradient autonomous ensemble: pval<0.001, size effect Z=19.602, n=124750.

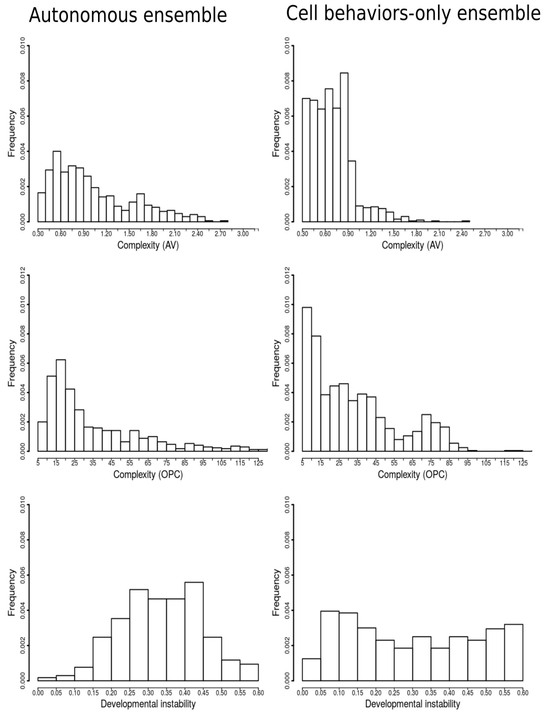


**Figure S11. Histograms of the autonomous ensemble and the cell-behaviors-only ensemble.** Both ensembles produce different distributions of complexity, but more importantly is to notice that the developmental instability histograms are not skewed to the left, as it does occur in the other ensembles. This indicates an increase in the average instability of the complex morphologies in the autonomous ensemble and in the cell-behaviors only ensemble.

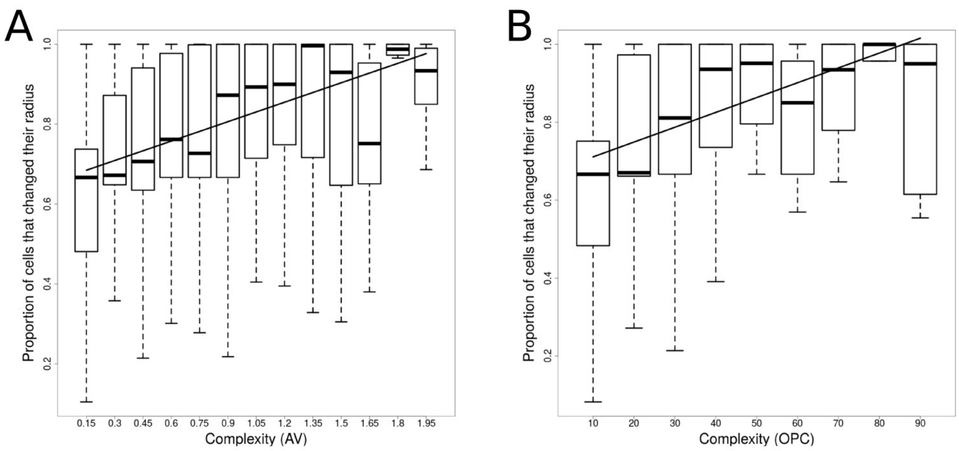


**Figure S12. Complexity requires changes in cell behaviors or node properties over extensive areas of the embryo.** The plots shows, per embryo, the proportion of cells that have changed their size ($p^{EQD}$) by more than 10% between the beginning and the end of development. Each embryo represents a point but we represent those by a boxplot. Outliers are not plotted, boxes enclose 50% of the data, the line in the box shows the median. The whiskers represent the range, which is 1.5 times the interquartile range. This effect is only evident in the case of the cell radius. A. uses angle variation as the complexity measurement (Spearman correlation: $r_s$=0.824 and pval=0.001) and B uses OPC as the complexity measurement (Spearman correlation: $r_s$=0.783 and pval=0.019). Spearman pval was calculated using a permutation test.

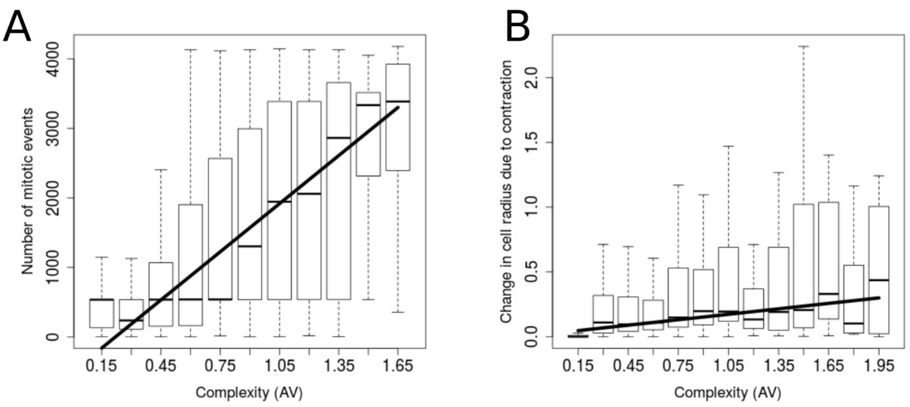


**Figure S13. Cell contraction and division were the cell behaviors most often associated with complex morphology**. Some cell behaviors have a clear effect on complexity. In **A** we show there is a positive correlation between cell division (number of mitotic events through a simulation) and AV complexity. In **B** we show there is a positive correlation between complexity and the change in average cell radius due to contraction. Cell radius is determined, in each iteration, by the sum of four components. Each of them is a node property that can change over simulation time, in **B** we show only the component due to active cell contraction. To plot the results, we construct boxplots with the individuals which fall in intervals of 0.15 AV complexity (x-axis). The line shows the linear regression using the median of each boxplot. **A** Spearman correlation: $r_s$=0.989 and pval<0.0001 and **B.** Spearman correlation: $r_s$=0.863 and pval=0.001. The pval for the Spearman correlation was calculated with a permutation test.

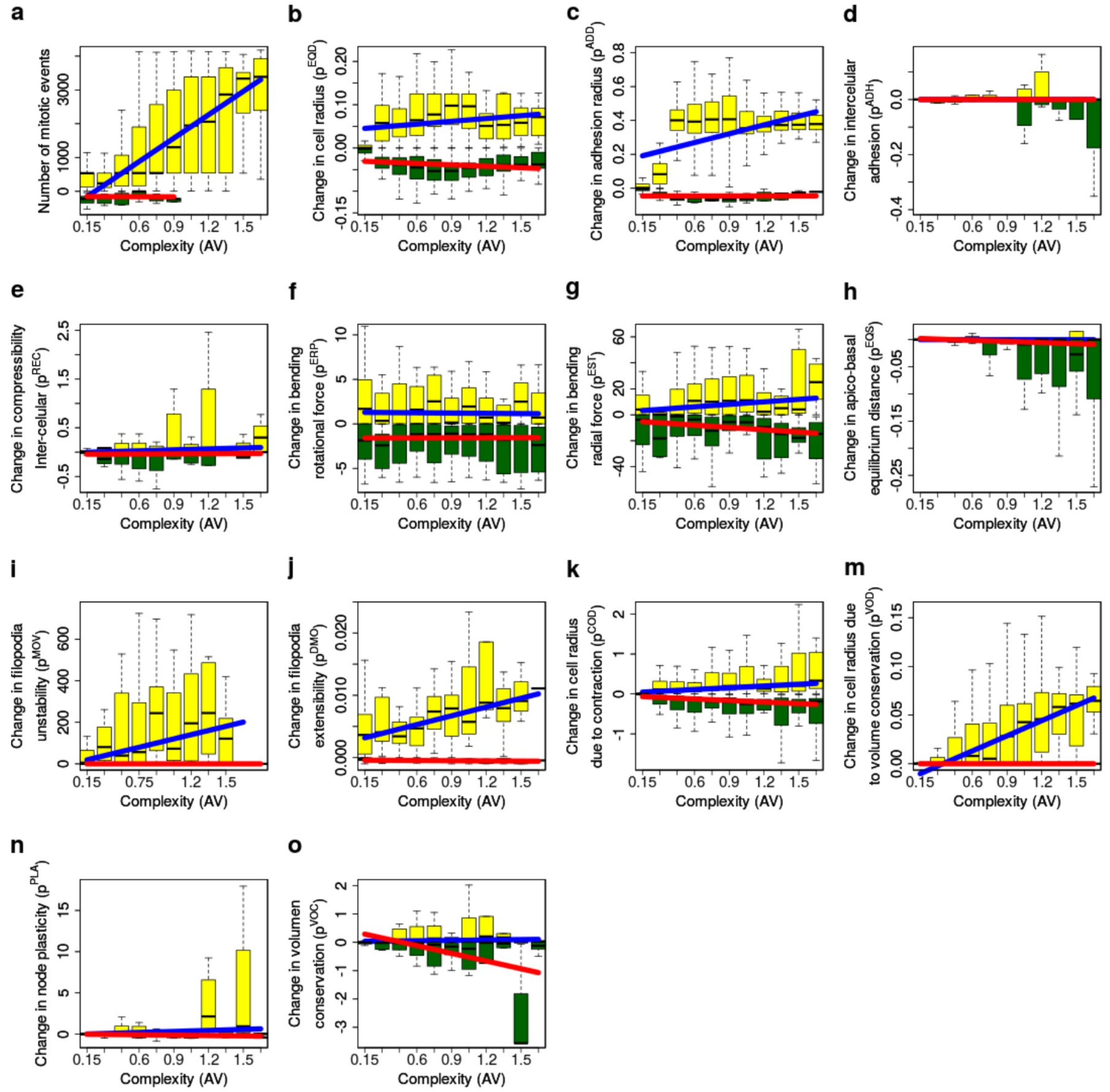

**Figure S14. Changes in node properties and cell behaviors versus complexity.** Some cell behaviors have a clear effect on complexity. In the ensemble changes in some cell behaviors and mechanical properties do not lead to enough viable (*e.g.* non-broken) embryos to build informative plots. The change in each node property or cell behavior is calculated as the difference in the average value of a node property across the embryo between the initial conditions and the final morphology. Regression lines are plotted separating simulations in which the value of a node property increases between the initial condition and the final morphology (blue line) and simulations in which it decreases (red line). The linear regression uses the median of each boxplot. Boxes enclose 50% of the data, the line in the box shows the median. The whiskers represent the range, which is 1.5 times the interquartile range. See table 1 table for the statistics of each correlation. See table 2 and (16) for detailed descriptions of each node property.

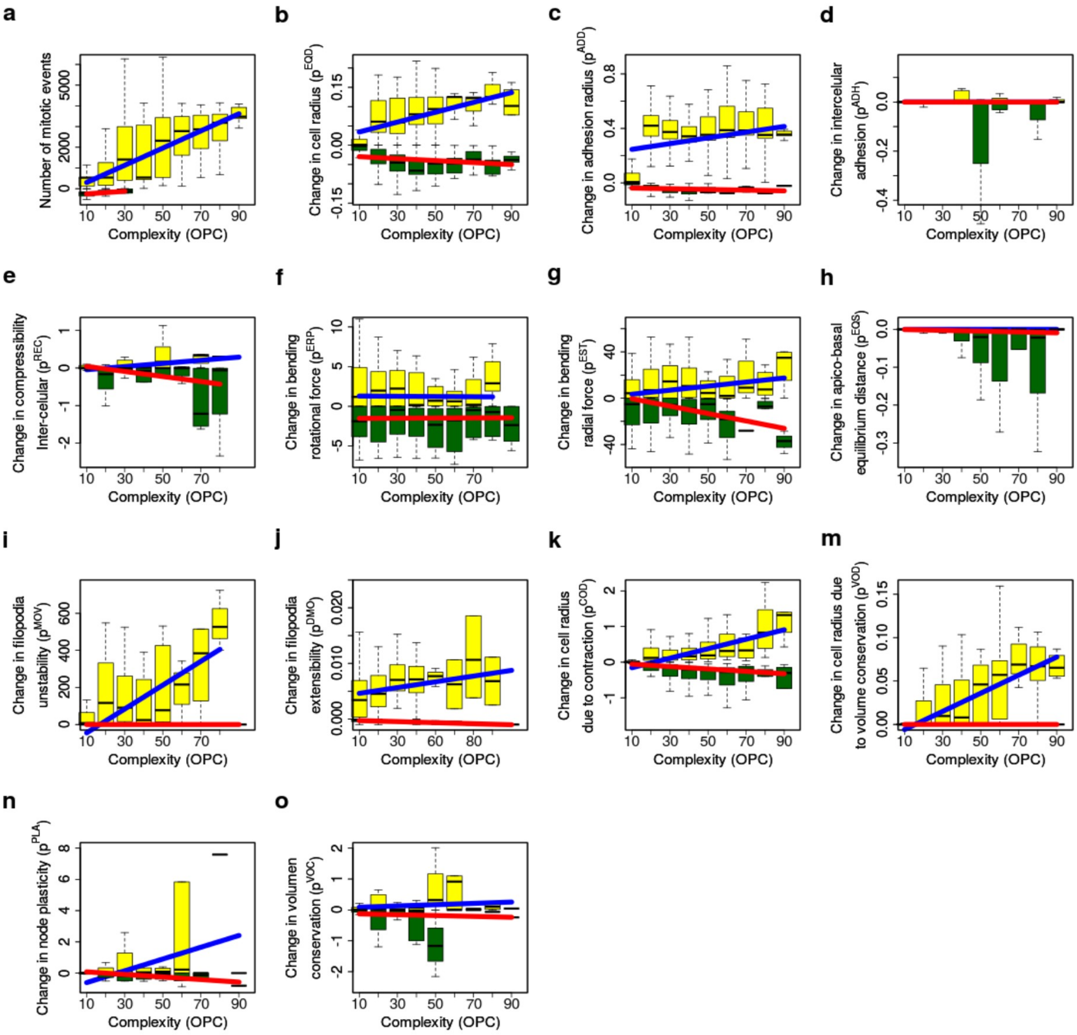


**Figure S15. As figure S13 but using OPC complexity.** See table 3 table for the statistics of each correlation.

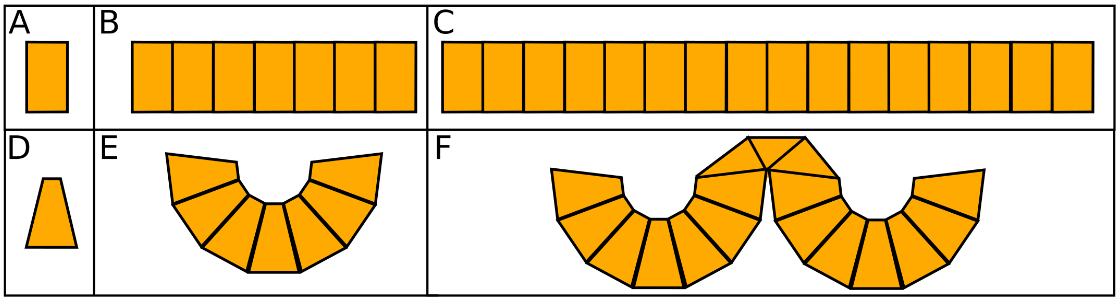


**Figure S16. Geometrical constraints in invagination.** In **A** we can see the shape of an individual cell that can be found in flat epithelia. **B** and C simply show two epithelia of different length. If due contraction the cell shape changes from that of **A** to that of **D**, the epithelial sheets will bend. If this occurs in a relatively small epithelium like the one in **B**, a single invagination as in **E** will form. Notice that the curvature of the invagination will depend of the shape of the individual cells, the more the constricted the cells, the higher the curvature will be. If this contraction occurs in a larger epithelium like the one shown in **C**, the epithelium does not bend into a single invagination because it is larger and due to geometry, only certain number of cells can fit into a single invagination. As a result two evaginations form, which have roughly the same number of cells as the invagination in **E**.

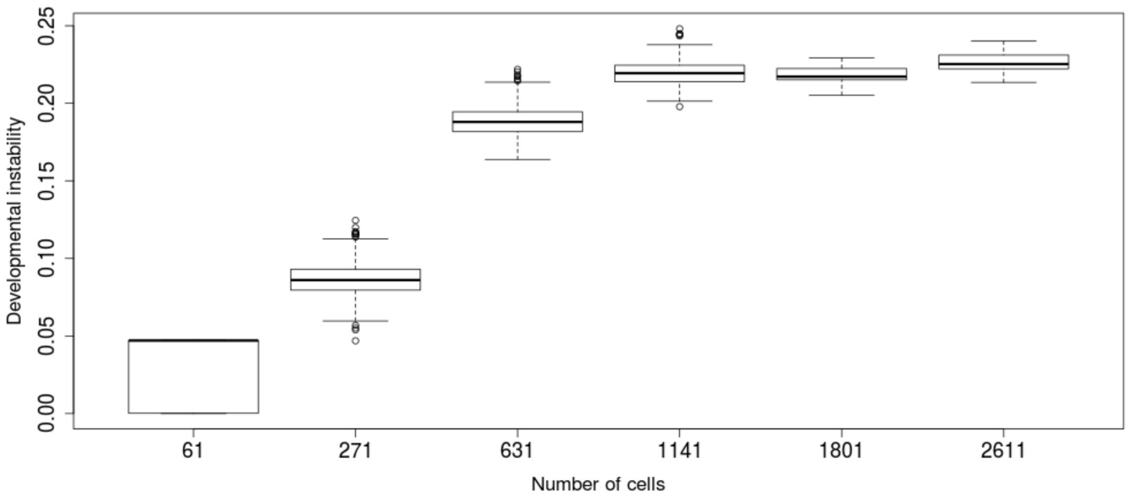


**Figure S17. The larger the number of cells contracting at the same time the larger the developmental instability**. To calculate developmental instability we simulate 30 times each initial developmental pattern shown in figure 4 with noise (changing the random seed). We then measure the EMD distance between all final morphologies. As we can see, developmental instability increases with embryonic size, although for these kind of morphology the EMD distances saturate at 1141 cells. Boxplots as in previous figures.

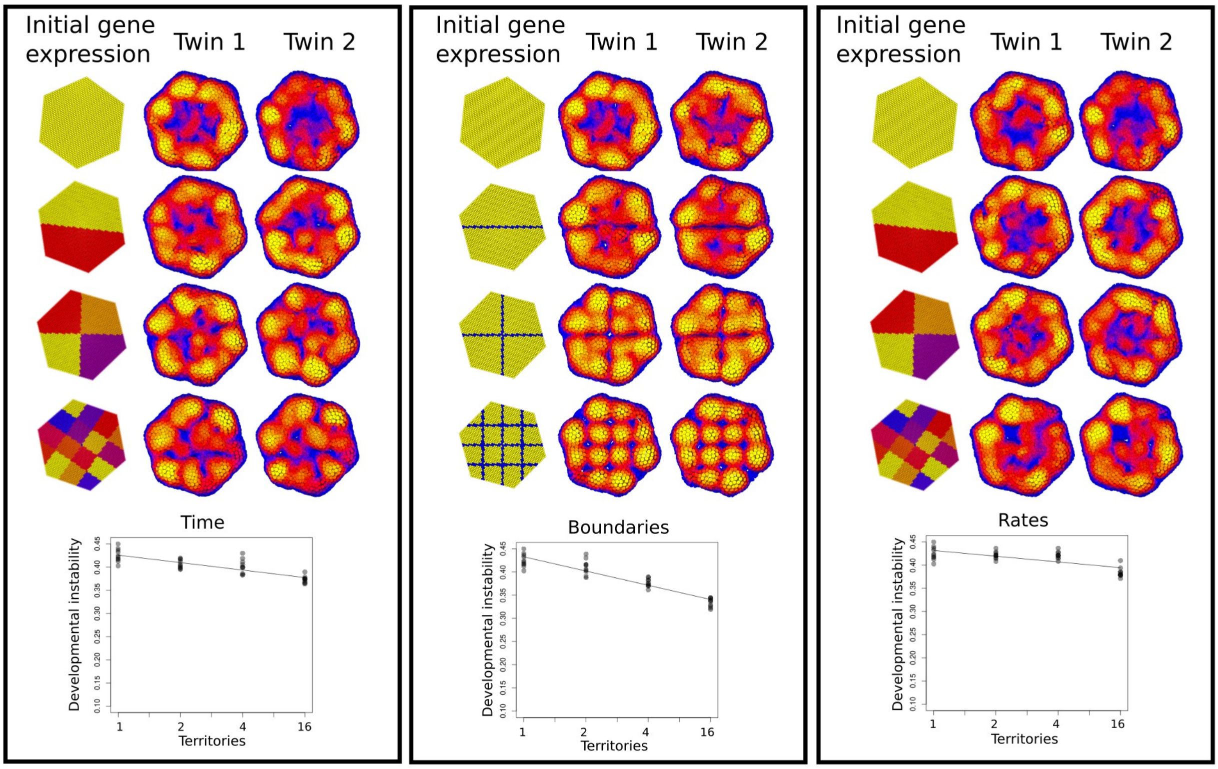


**Figure S18. Splitting the epithelium into regions of gene activity that regulate cell apical contraction can increase the developmental stability.** In each column we show the initial gene expression on the left (colors shows the genes being express) and the two of the final morphologies on the right, each of them we call a twin (color shows the cell position in the z-axis). The first column shows the results of splitting the embryo into regions that contract at different times (every 0.5 model time units), each of this regions will start contracting at different time points, but they all contract in the same way once they start contracting. The middle column show the results of splitting the embryo into territories of contracting cells (yellow cells) with non-contracting boundaries (blue cells). All the contracting cells contract at the same time and in the same way. The right column show the results of splitting the embryo into territories of cells that contract at different rates (although the variance was keep the same between the simulations of different territories). It is visually clear that splitting the embryos into territories increases the stability, specially when paying attention to the location of ridges (in yellow) and bumps (in blue). At the bottom of each column we show the results of quantifying the results. To calculate developmental instability we simulate 5 times each initial conditions with noise (changing the random seed). As before, developmental instability is measured by calculating the EMD distances between all the twins. Each point in the plot represent one of this distances. The intensity of the gray of the points indicates how many overlapping points there are. The line shows the linear regression. Time: Spearman correlation: $r_s$=-0.811, pval<0.001 . Boundaries: Spearman correlation: $r_s$=-0.922, pval<0.001. Rate: Spearman correlation: $r_s$=-0.622, pval<0.001. The pval for the Spearman correlation was calculated with a permutation test.

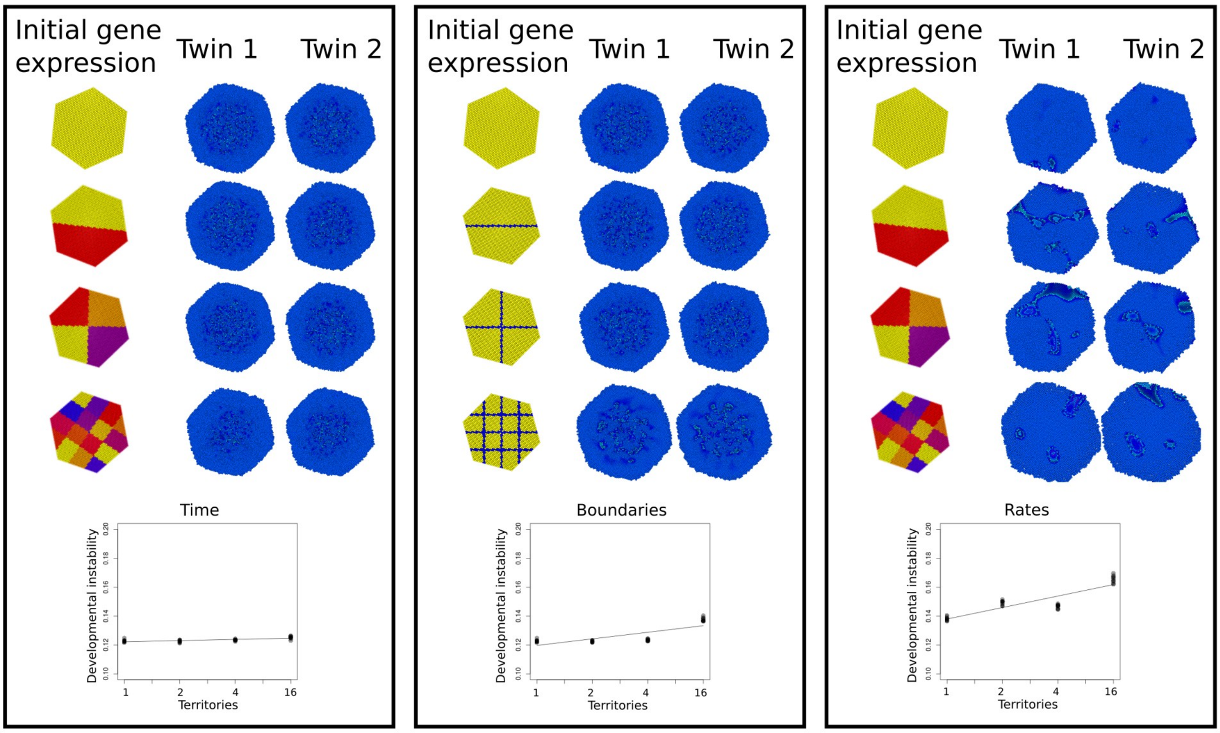


**Figure S19. Splitting the epithelium into regions of gene activity that regulate cell division does not have a clear effect on developmental stability.** As in figure S17 but genes regulate cell division. The results for the quantification are: Time: Spearman correlation: $r_s$=0.726, pval<0.001. Boundaries: Spearman correlation: $r_s$=0.748, pval<0.001. Rate: Spearman correlation: $r_s$=0.794, pval<0.001.

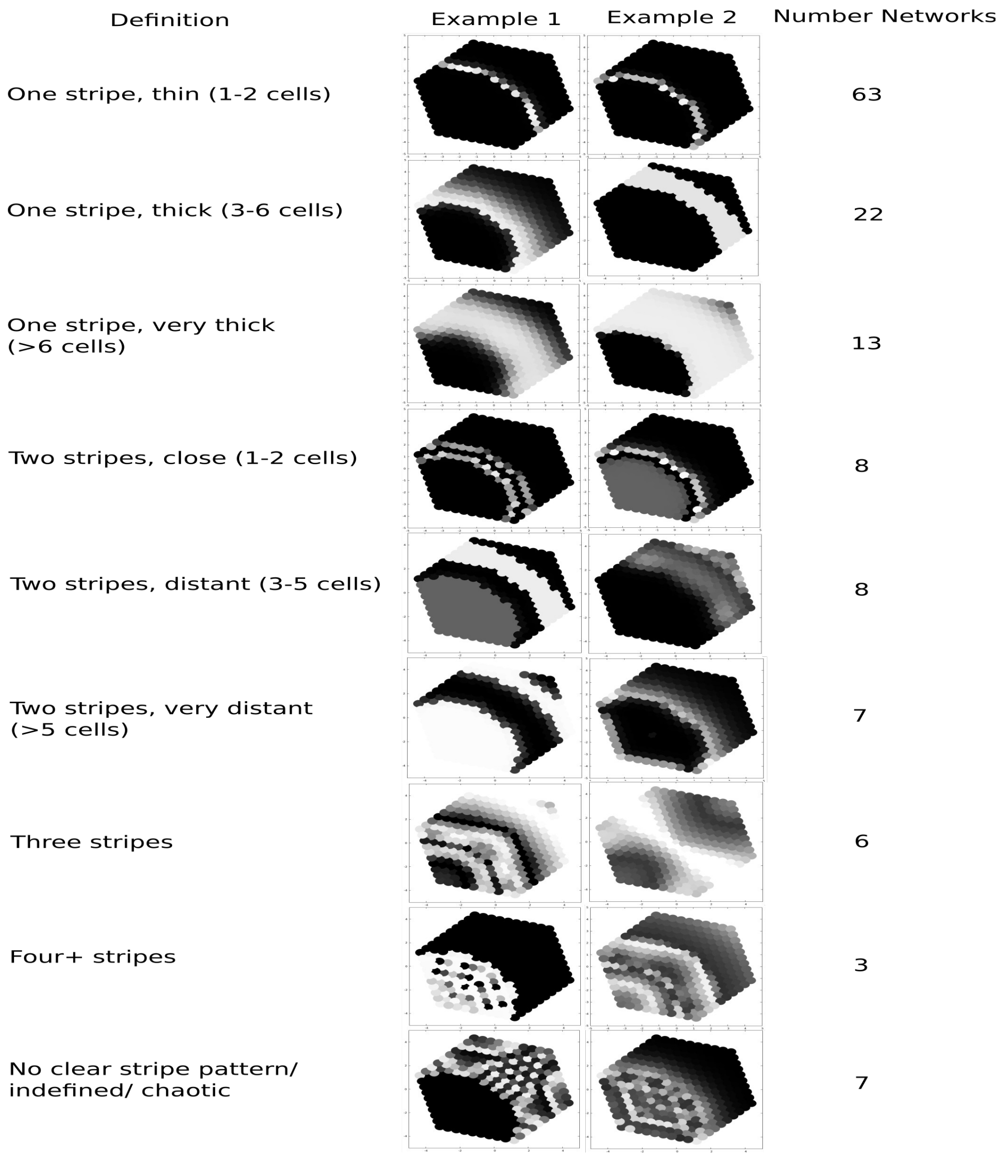


**Figure S20. Classification of signaling ensemble.** The gene networks from the signaling ensemble were classified depending on the gene expression patterns they produced. Only gene networks that produced stable and heterogeneous patterns were classified. The left column includes the classification arguments. In the middle we can see two examples for each of the categories. On the right we can see the number of gene networks that were found for each of the categories.

### Tables

Table 1. **Statistics of spearman correlarions calculated from figure S14.**

| | | Increasing | | Decreasing | |
|---|---|---|---|---|---|
| | | pval | $r_s$ | pval | $r_s$ |
| a | Number of mitotic events | <0.001 | 0.988 | 0.708 | -0.200 |
| b | Change in cell radius | 0.306 | 0.345 | 0.524 | -0.218 |

| | | | | | |
|---|---|---|---|---|---|
| c | Change in adhesion radius | 0.613 | 0.172 | 0.675 | 0.145 |
| d | Change in intercelular adhesion | 0.311 | 0.336 | <0.001 | -0.882 |
| e | Change in comprensibility (inter-cellular) | 0.942 | -0.027 | 0.487 | 0.236 |
| f | Change in bending rotational force | 0.738 | -0.118 | 0.903 | 0.045 |
| g | Change in bending radial force | 0.219 | 0.4 | 0.156 | -0.463 |
| h | Change in epithelial spring elastic force | 0.099 | 0.527 | 0.001 | -0.864 |
| i | Change in filopodia unstability | 0.037 | 0.716 | <0.001 | -0.933 |
| j | Change in filopodia extensibility | <0.001 | 0.963 | 0.348 | -0.312 |
| k | Change in cell radius due to contraction | 0.001 | 0.863 | 0.014 | -0.745 |
| l | Change in cell radius due to volumen conservation | <0.001 | 0.991 | 0.072 | 0.843 |
| m | Change in cell plasticity | 0.965 | -0.018 | 0.218 | -0.4 |
| n | Change in volumen conservation | 0.289 | 0.354 | 0.037 | -0.636 |

Gray boxes show p-values < 0.05.

Table 2. Summary of node properties

| |
|---|
| Common to all types of node |

| Name | Symbol | Description |
|---|---|---|
| Intercellular adhesion | $p^{ADH}$ | Adhesion force between nodes |
| Cell compressibility | $p^{REC}$ | Strength of repulsion force between nodes |
| Filopodia extensibility | $p^{DMO}$ | Nodes that move due to noise, do so in a random direction for a random distance between 0 and $p^{DMO}$ |
| Filopodia unstability | $p^{MOV}$ | Probability of accepting a node movement even when its new position has higher potential energy than its position before movement. |
| Node plasticity | $p^{PLA}$ | Specifies how plastic a node is, i.e., how the nodes $p^{PLD}$ will change due to pressure |
| Degree of differentiation | $p^{DIF}$ | Determines how differentiated a node is. As differentiation increases, changes in nodes slow down |
| Equilibrium radius | $p^{EQD}$ | Distance at which nodes start repelling each other |
| Contraction component of $p^{EQD}$ | $p^{COD}$ | Component of node's size, $p^{EQD}$ , due to cell contraction or expansion. |
| Growth component of $p^{EQD}$ | $p^{GRD}$ | Component of node's size, $p^{EQD}$ , due to growth or apoptosis. |
| Plasticity component of $p^{EQD}$ | $p^{PLD}$ | Component of node's size, $p^{EQD}$ , due to plasticity |
| Adhesion radius | $p^{ADD}$ | Distance at which nodes start to adhere to each other |
| Amount of stored ECM | $p^{ECM}$ | Cells that produce ECM will accumulate it before secreting it. Once $p^{ECM}$ reaches the value of the model parameter $M_{ECM}$, a node is secreted and $p^{ECM}$ is set back to 0. |
| Only for epithelial nodes | | |
| Rotation force component resistance | $p^{ERP}$ | Weight of the rnon-adial component of the epithelial rotation force. This force rotates the cylinders until the their apical-basal vector is normal to the surface plane at that position |
| Radial force component resistance | $p^{EST}$ | Weight of the radial component of the epithelial rotation force. Radial force reduces sliding from apical or basal nodes along the apical-basal direction from neighbor cylinders |
| Apico-basal equilibrium distance | $p^{EQS}$ | Equilibrium length of the spring between both nodes in a cylinder |
| Volume conservation component of $p^{EQD}$ | $p^{VOD}$ | Component of node's size, $p^{EQD}$ , due to cell volume conservation |

Table 3. Statistics of the linear regressions calculated from figure S15.

| | | Increasing | | Decreasing | |
|---|---|---|---|---|---|
| | | pval | $r_s$ | pval | $r_s$ |
| a | Number of mitotic events | <0.001 | 0.983 | 0.330 | 1 |
| b | Change in cell radius | 0.003 | 0.866 | 0.547 | -0.233 |
| c | Change in adhesion radius | 0.1 | 0.862 | 0.467 | -0.283 |
| d | Change in intercelular | 0.433 | 0.3 | 0.061 | -0.667 |

| | | | | | |
|---|---|---|---|---|---|
| | adhesion | | | | |
| e | Change in comprensibility (inter-cellular) | 0.084 | 0.667 | 0.461 | -0.309 |
| f | Change in bending rotational force | 0.841 | -0.095 | 0.748 | -0.133 |
| g | Change in bending radial force | 0.360 | 0.35 | 0.003 | -0.750 |
| h | Change in epithelial spring elastic force | 0.043 | 0.700 | 0.148 | -0.533 |
| i | Change in filopodia unstability | 0.028 | 0.786 | 0.007 | -0.833 |
| j | Change in filopodia extensibility | 0.171 | 0.547 | 0.001 | -0.899 |
| k | Change in cell radius due to contraction | 0.003 | 0.728 | <0.001 | 0.844 |
| l | Change in cell radius due to volumen conservation | 0.001 | 0.916 | 0.439 | -0.3 |
| m | Change in cell plasticity | 0.801 | 0.100 | 0.003 | -0.867 |
| n | Change in volumen conservation | 0.383 | 0.383 | 0.708 | -0.15 |

Gray boxes show p-values < 0.05.

Table 4. Limits of the parameters used in the model.

| **Ranges of the model parameters** | | | |
|---|---|---|---|
| Parameter | Symbol | Minimum | Maximum |
| **Molecular parameters** | | | |
| Trancription | $t$ | 0 | 31.62 |
| Degradation rate | $\mu$ | 1 | 32 |
| Diffusion | $D$ | 0.0021 | 0.21 |

| | | | |
|---|---|---|---|
| **Node properties** | | | |
| See also 2.7.4 | | | |
| Components of $p^{EQD}$ | $p^{COD}$, $p^{GRD}$, $p^{PLD}$, $p^{VOD}$ | 0.0083 | 0.83 |
| Adhesion radius | $p^{ADD}$ | 0.00375 | 3.75 |
| Intracellular plasticity | $p^{YOU}$ | 0.00053 | 5.3 |
| Cell compressibility to nodes from a different cell | $p^{REC}$ | 0.00053 | 5.3 |
| Apico-basal equilibrium distance | $p^{EQS}$ | 0.0083 | 0.83 |
| Filopodia unstability | $p^{MOV}$ | 0.0001 | 10 |
| Filiopodia extensibility | $p^{DMO}$ | 0.075 | 7.5 |
| Degree of diferentation | $p^{DIF}$ | 0.075 | 0.1 |
| **Cell behaviors** | | | |
| Cell division | $C_k$ | 0.02 | 1 |
| Apoptosis | $C_{APO}$ | 0 | 0.005 |
| Epithelial mesenchymal transition | $C_{EMT}$ | 0 | 0.05 |
| ECM secretion | $C_{ECM}$ | 0 | 0.05 |

Table 5. Parameters and variables of the initial conditions.

| Parameters and variables value at the initial conditions | | |
|---|---|---|
| **Global model parameters** | | |
| Temperature | $M_{TEM}$ | 0.001 |
| Maximal compression allowed in a cell to allow growth in it | $M_{MCO}$ | -0.1 |
| Maximum node length of movement | $M_{RMA}$ | 0.001 |
| Maximal number of nodes allowed | $M_{MAN}$ | 5000 |
| Time a node can be alone before dying | $M_{TAL}$ | 10 |
| Minimal $p^{EQD}$ allowed | $M_{EMI}$ | 0.0083 |
| Maximal $p^{EQD}$ allowed | $M_{EMA}$ | 0.83 |
| Amount of extra-cellular matrix that has to accumulate in a node before an ECM node is secreted | $M_{ECM}$ | 0.25 |
| Maximum $\delta$ | $M_{DMA}$ | 0.01 |
| Minimum $\delta$ | $M_{MDI}$ | 0.0001 |
| Proportion of nodes subject to | $M_{NOI}$ | 0.001 |

| | | |
|---|---|---|
| noise per iteration | | |
| Maximal number of nodes that can interact with a node | $M_{MNN}$ | 500 |
| Accuracy of the numerical integration | $M_{DDA}$ | 0.001 |
| diffusion coefficient of $p^{EQD}$ | $M_{DID}$ | 1 |
| Sets the size of the exclusion sphere when using the Gabriel method | $M_{GAB}$ | 0.85 |
| Maximum $p^{EQD}$ allowed for an epithelial node due to deformation | $M_{DFE}$ | 0.5 |
| **Model variables** | | |
| Equilibrium radius | $p^{EQD}$ | 0.25 |
| Adhesion radius | $p^{ADD}$ | 0.35 |
| Intercellular adhesion | $p^{ADH}$ | 10 |
| Cell compressibility to nodes from a different cell | $p^{REC}$ | 10 |
| Rotation force component resistance | $p^{ERP}$ | 4.8 |
| Radial force component resistance | $p^{EST}$ | 62.5 |
| Apico-basal equilibrium distance | $p^{EQS}$ | 0.5 |
| Filopodia unstability | $p^{MOV}$ | 0.0001 |
| Filopodia extensibility | $p^{DMO}$ | 0.01 |
| Amount of stored ECM | $p^{ECM}$ | 0 |
| Contraction component of $p^{EQD}$ | $p^{COD}$ | 0 |
| Growth component of $p^{EQD}$ | $p^{GRD}$ | 0.25 |
| Plasticity component of $p^{EQD}$ | $p^{PLD}$ | 0 |
| Degree of differentiation | $p^{DIF}$ | 0 |
| Node plasticity | $p^{PLA}$ | 1 |
| Volume conservation component of $p^{EQD}$ | $p^{VOD}$ | 10 |
